# Systems-level Plant Responses Reveal *Pseudomonas*-Mediated Growth Promotion in *Brachypodium* Under Nitrogen Limitation

**DOI:** 10.1101/2025.11.03.686263

**Authors:** S. Sanow, M. Mantz, H. Wissel, A. Bhatnagar, E. Sturm, J. M. Kelm, R. Walker, A. Lücke, G. Schaaf, P. Huesgen, U. Roessner, M. Watt, B. Arsova

## Abstract

Plant molecular adaptation to plant growth promoting bacteria (PGPB) under nutrient stress remains unclear, yet is essential for advancing PGPB use in agriculture. The model grass *Brachypodium dystachion* was studied together with *Pseudomonas koreeensis (Pk)* at two nitrogen (N) conditions. Non-invasive shoot phenotyping showed an immediate response to low-N, while beneficial effects of *Pk* became quantifiable after day 19. Increased N content in inoculated plants, along with *Pk’s* ability to grow on N-free media, suggests bacterial N contribution at deficient N. In low-N conditions, *Pk*-inoculated plants showed 33.2% more N than uninoculated controls and biomass comparable to high-N plants. *Pk* had no effect under sufficient N. Proteomics and lipidomics revealed that lipid profiles were primarily shaped by N availability, while protein abundance responded to both *Pk* and N status. Inoculated low-N plants displayed protein profiles resembling those of high-N controls, with some distinct exceptions. The plant-microbe interaction is dynamic and developed over 3 weeks, leading to increased biomass and N content. Root proteins strongly induced by *Pk* under low-N included lipid degradation enzymes, N transporters, and regulatory proteins, suggesting a coordinated remodelling of energy metabolism supporting whole-plant biomass and increased abundance of N uptake proteins.

## Introduction

Agriculture today requires biological alternatives that can reduce mineral fertiliser input, to decrease environmental pollution, while maintaining global food security. One promising strategy involves the use of plant growth-promoting bacteria (PGPB), which can either enhance resource availability or modify plant physiology to improve nutrient foraging — or both. However, their inconsistent performance across environmental conditions remains a major limitation (Lopes et al., 2021). One way to address this inconsistency is by dissecting the plant response (or adaptation) to the presence of a potential PGPB under varying abiotic conditions. Kuang *et al*. (2022), demonstrated that Brachypodium plants inoculated with *Herbaspirillum seropedicae* exhibited different modes of growth promotion under two nitrogen (N) conditions. The study reported differences in N uptake rates (and N-forms), transcript regulation, and temporal responses between the two N conditions (Kuang *et al*., 2022). However, the systemic response to the presence of a beneficial bacterium is still not fully understood. lant–microbe interactions trigger complex signalling cascades involving proteins, lipids, and metabolites. An article by Schillaci *et al*., 2021 highlighted the role of lipid biomolecules in plant-microbe interaction. We are still missing an integrated dataset across several levels of metabolites- to begin to comprehend the plasticity of the plant molecular response.

The genus *Pseudomonas* is ubiquitous throughout the globe and encompasses bacteria that can be pathogenic, saprophytic, or beneficial to the plant (Preston 2004; Volke & Nikel 2020; Rooney et al., 2020). Among them, we focus on species which improve plant performance at sub-optimal N conditions (Wang et al., 2020; Sanow et al., 2023; Singh et al., 2023). *Pseudomonas spp*. have several mechanisms for beneficial interaction with plants, including the increase of plant N-content through: (i) Ammonification and denitrification, (ii) production of secondary metabolites to stimulate N-fixation in adjacent bacterial strains, and (iii) Biological Nitrogen Fixation (Sah & Singh 2021; Tu et al., 2021; Sanow *et al*., 2023).

*Pseudomonas koreensis* Ps 9-14T was first described in 2003 as a new species from Korean soils (Kwon et al., 2003), which was reported to increase the abundance of other microbes to increase protection against pine wilt disease (Han et al., 2021). The strain *Pseudomonas koreensis* IB-4 was described in 2016 as a potential new strain for plant biocontrol in potato against pathogenic fungi *Fusarium, Bipolaris*, and *Alternaria* (Rafikova et al., 2016). Notably, genetic analysis revealed that *P. koreensis* (hereafter *Pk*) harbors genes associated with atmospheric nitrogen fixation. Nitrogenase activity in *Pk* was confirmed using the ethylene reduction assay (Rafikova et al., 2016). However, the ability of *Pk* to increase nitrogen content *in planta* has not been demonstrated.

Given this potential, it is worth investigating whether *Pk* can serve as an effective PGPB in the context of reducing synthetic nitrogen fertilizer use.

Ammonium (NH_4_^+^) and Nitrate (NO_3_^-^) are the main uptake forms of N into the plant, conducted by the Ammonium Transport (AMT) family and Nitrate Transport Family (NRT) Feng *et al*., 2020. NH_4_^+^ and NO_3_^-^ are then assimilated via several proteins of the central N metabolism, including nitrate reductases (NR), nitrite reductases (NiR), glutamine synthetases (GLN/GS), and glutamate synthase (GOGAT) (Masclaux-Daubresse, *et al*. 2010). The aforementioned proteins are crucial targets to study N uptake and assimilation under N-fixing conditions.

In this study, we investigated the response of Brachypodium to *Pseudomonas koreensis* Ps 9-14T inoculation under two N conditions, by integrating time-resolved data of phenotype, elemental analyses, proteomics and lipidomicsProteomic profiling at 21 days post-inoculation provided insights into whole-plant protein abundance, while lipidomics enabled us to examine dynamic lipid profiles — still a largely unexplored dimension in plant–microbe interactions (Macabuhay et al., 2022). As such, *Pseudomonas koreensis* Ps 9-14T was used (i) as a bacterial trigger to study plant adaptation processes to the presence to a PGPB, and (ii) as a candidate beneficial microbe to evaluate its potential to improve growth in grass species, which include many globally important crops.

## Material & Methods

### Plant cultivation system preparation

*Brachypodium distachyon* Bd21-3 seeds were surface sterilized (Sasse et al., 2019), dehusked, and sown into modified Plant Cultivation Vessels (PCVs; GA-3, Kisker Biotech; Supplemental Figure 1) filled with 900 g of 0.7–1.4 mm quartz sand. Containers (except connectors) were autoclaved and oven dried. 54 g of modified Hoagland solution (1.76 ml 0.5 M KH_2_PO_4_, 2.64 ml 1M MES (pH 5.7), 4.39 ml 10 mM FeSO_4_ + 10 mM Na_2_EDTA, 1.76 ml 1 M CaCl_2_, 1.76 ml 0.375 M MgSO_4_, 1.76 / 17.6 ml 1 M NH_4_NO_3_ (low / high N) and 0.44 ml micronutrients (50 mM H_3_BO_3_, 10 mM MnSO_4_, 2 mM ZnSO_4_, 5 mM KI, 0.2 mM Na_2_MoO_4_, 0.1 mM CoCl_2_, 1 mM CuSO_4_) in a total volume of 1L) was added to each under sterile conditions. Three sowing holes were made using a template. (Supplemental figure 2).

### Seed sterilization, sowing and stratification

*Brachypodium distachyon* Bd21-3 (Brachypodium) (Vogel & Hill, 2007) seeds were dehusked and surface sterilized as described in Sasse et al., 2019. A single seed was placed in each of the three pre-made holes with sterile tweezers and covered with sand. At 5 DAS, excess seedlings were removed to ensure homogenous shoot height.

### Bacterial cultivation and inoculation

*Pseudomonas koreensis* Ps9-14T (*Pk*) (DSM16610, Germany) was cultivated using Luria-Bertani (LB) medium with optional 1% agar (w/v) for plates at 29 °C, 220 rpm (liquid culture). A pre-recorded bacterial growth curve was established with an OD_600_ of 0.5 corresponding to 1x10^8^ CFU ml^-1^. Bacteria from the overnight culture were centrifuged at 3000 rpm for 15 minutes and the bacterial pellet was resuspended in 1 ml of the respective Hoagland solution and further diluted to 5x10^6^ CFU ml^-1^. Each seedling received 100 µl inoculum (5 × 10⁵ CFU); controls received media. PCV were imaged for non-invasive shoot phenotyping (c.f. Non-invasive shoot phenotyping) before being closed and transferred to growth chambers.

### Plant Growth Conditions and experimental setup

Plants were grown in growth chambers (Percival Scientific, Model AR-22L) with 16/8 h day/night period, at 23/18 °C with 100-140 μmol m^-2^s^-1^ light and 35 - 40% relative humidity. PCVs were placed in three identical chambers and randomized per imaging time-point.

In three independent experiments plants were cultivated for 21 DAS and were removed from the growth chamber at 5 DAS for inoculation, non-invasive shoot phenotyping (every 2 days), and watering at 7 and 14 DAS (Supplemental Fig. 3).

### Non-invasive shoot phenotyping

Shoots were imaged from three angles using a smartphone (Samsung Note8) in a beige-background photobox inside a sterile hood (Supplemental Fig. 4). Projected leaf area was calculated via HSV color segmentation (Müller-Linow et al., 2022) with thresholds: Hue 44.4– 127.5, Saturation 66.6–255, Value 6.75–255. The shoot phenotyping interval was: 5, 7, 9, 11, 13, 15, 17, 19, 20, 21 DAS. Invasive harvests were conducted at 19, 20, 21 DAS (Supplemental Fig. 3).

### Invasive root and shoot phenotyping

A total of 44 plants were harvested at 19, 20, 21 DAS. Five plants at each time point and treatment were used for invasive phenotyping and 6 plants for molecular analyses.

Roots were washed with sterile Milli-Q water and separated from shoots. After recording fresh weight, tissues were either frozen in liquid N₂ or scanned (WinRhizo, Regent Instruments) for morphological analysis. Samples were oven-dried (65 °C, 3 d), weighed, and homogenized (Retsch MM400, 3 mm bead, 30 Hz, 1 min) for C/N analysis (Cario EL Cube, Elementar) with thermal conductivity detection. Root images were analyzed using PaintRhizo (Nagel et al., 2012) to determine primary and lateral root length

The WinRhizo generated root images were analysed using the software package PaintRhizo (Nagel et al., 2012) and the total root length was dissected into primary seminal root length and lateral root length.

### Data Analysis and Statistics

Statistical analyses were performed using two-way ANOVA with a confidence level of 95% followed by a post-hoc Tukey Honestly Significant Difference (HSD) test (FDR < 0.05). Correlations were calculated using the Pearson coefficient.

### Sample preparation for protein extraction

Roots and shoots from two plants were pooled per biological replicate (≥100 mg roots; ∼55 mg shoots). From each, 30–50 mg was homogenized (ball mill, 5 × 1 min) without thawing. Proteins were extracted with PBS + 2% SDS (100 µl per 10 mg fresh weight) and quantified via BCA assay. For digestion, 25 µg protein was denatured (10 mM DTT, 90 °C, 15 min), alkylated (50 mM chloroacetamide, 30 min dark, RT), and quenched (50 mM DTT, 20 min). SDS removal and trypsin digestion (1:100) followed the SP3 protocol (Demir et al., 2022), with overnight incubation at 37 °C. Peptides were cleaned using modified SDB-RP StageTips (Rappsilber et al., 2003), eluted, vacuum-concentrated, and resuspended in 0.1% formic acid. 2 μl of the sample were used to determine peptide concentration at the Nanodrop, before injection of 1 µg peptides into LC-MS/MS.

### Proteomics data acquisition

1 µg peptides per sample was analyzed via LC-MS/MS using an UltiMate 3000 RSLC nano-HPLC system (Thermo) with μPAC pillar array trap and analytical columns (1 cm and 50 cm flowpath, respectively, PharmaFluidics) on-line coupled to an Impact II Q-TOF mass spectrometer (Bruker) with CaptiveSpray source (Bruker) as described (Beck et al., 2015). Peptides were separated using an 80 min 5–32.5% ACN gradient (600 nl/min, 40 °C) and tandem mass spectrometry data were acquired using a data independent acquisition (DIA) scheme with MS1 survey scans from 400 to 1200 m/z followed by acquisition of MS2 fragment ion in 32 selection windows of 25 m/z with 0.5 m/z overlap, scanning a mass range from 200 to 1750 m/z at 15 Hz.

### Proteomics data analysis and visualization

Peptides were identified by matching the acquired data to a predicted spectral library using DIA-NN version 1.8 (Demichev et al. 2020) with the Brachypodium sequences in the UniProt database (UP000008810_15368, release 2022_04). Parameters were set to trypsin as digestion enzyme with up to two maximum missed cleavages, a maximum of two variable peptide modifications (variable Met oxidation), a 1% Precursor FDR and fixed Cys carbamidomethylation. Peptide lengths of 7 to 30 amino acids and a precursor charge range of two to four were considered. The query was set in double-pass mode with match between runs (MBR) enabled and heuristic protein interference disabled. Mercator4 version 2.0 (Schwacke et al., 2019) was used for functional annotation of the protein sequences (supporting data).

Data were further processed with the Perseus software package v1.6.15.0 (PMID: 27348712). Comparative analysis considered only proteins quantified in all replicates of one condition (100% valid values), with significant differences assessed by ANOVA (FDR < 0.05 and post-hoc Tukey HSD <0.02). Differential proteins were visualized with MapMan v3.5.1R2 on modified N-metabolism maps (Wang et al., 2012; Arsova et al., 2012; Feng et al., 2020). The images for kinase families and lipid biosynthesis were used directly from the MapMan website (https://mapman.gabipd.org/web/guest/mapman-download).

### Confirmation of plant inoculation

*Pk* identification was confirmed via MaxQuant search against *P. koreensis* (UP001139955, 2023_04) with identical settings.

### LC-MS untargeted analysis for lipids

Root tissues (∼20 mg) were extracted following Kehelpannala et al. (2019) and dried in vacuo (Christ Alpha 2-4 LSCplus). Lipids were reconstituted in 100 µl butanol:methanol (1:1 v/v, 1% ammonium acetate), vortexed, shaken (20 °C, 1400 rpm, 10 min), and centrifuged (RT, 5 min, 13,000 rpm). 7 μl of each sample were aliquoted and pooled together to prepare pooled biological quality control (PBQC) sample. This PBQC sample was injected onto the column periodically during the LC-MS batch acquisition to monitor the performance and any variability caused due to the instrument.

Samples were acquired on a high-resolution Q-TOF UHPLC mass spectrometer (Sciex TripleTOF 6600) interfaced with ExionLC ultra-high pressure liquid chromatography (UHPLC). Data were acquired using Sequential Window Acquisition of all Theoretical Mass Spectra (SWATH-MS) mode. The total runtime for each sample acquisition was 30 min with a flow rate of 0.26 ml/min. Gradient conditions followed Kehelpannala et al. (2019). Source settings: curtain gas 35, gas 1 & 2 at 25, temp 250 °C, voltage 4500 V. Mass range: TOF MS 100– 1700 m/z; SWATH MS/MS 100–1700 m/z with 16.2 Da windows. Data was processed using MSDIAL ver.5.1.230719 (Tsugawa et al., 2015). The MS data presented corresponds to six individual biological replicates for each treatment group per day. Data are available via MetaboLights.

### Growth of bacteria on nitrogen-free media

N-fixation ability was tested using nitrogen-deficient malate medium (NFM; Weselowski et al., 2016). *Pseudomonas koreensis Ps 9-14T*, *P. syringae pv lapsa*, *P. taiwanensis VLB120*, and *E. coli* DH5α were pre-cultured in LB (28 °C, 220 rpm o/N). Bacteria were spread on LB and nitrogen-deficient malate medium (NFM) supplemented with 50 mg/L yeast extract (NFM+) plates. After one day of incubation, colonies were spread on LB and NDM plates, followed by incubation for 3 days at 28 °C and 36 °C for Pseudomonas and E. coli, respectively.

## Results

Three independent plant experiments, (Supplemental Fig. 3) were conducted with four conditions: Low-N, Low-N + *Pk*, High-N and High-N + *Pk*. The results were reproducible as shown with the correlation coefficients among the experiments (r = 0.98, 0.98, 0.99; Supplemental Fig. 5).

### *Pk* inoculated Brachypodium under limited N shows faster growth and higher biomass

Non-invasive measurement of the leaf area revealed that up to 15 DAS, plants mainly responded to the different N availability. At 17 DAS the inoculated plants at low-N started showing an increase in the leaf area. At 20 and 21 DAS this was 26.7% and 39.9%, respectively, to their non-inoculated low-N controls (Fig. 1 a). *Pk* inoculation did not impact the leaf area at sufficient N (Fig. 1 a).

**Fig. 1.**
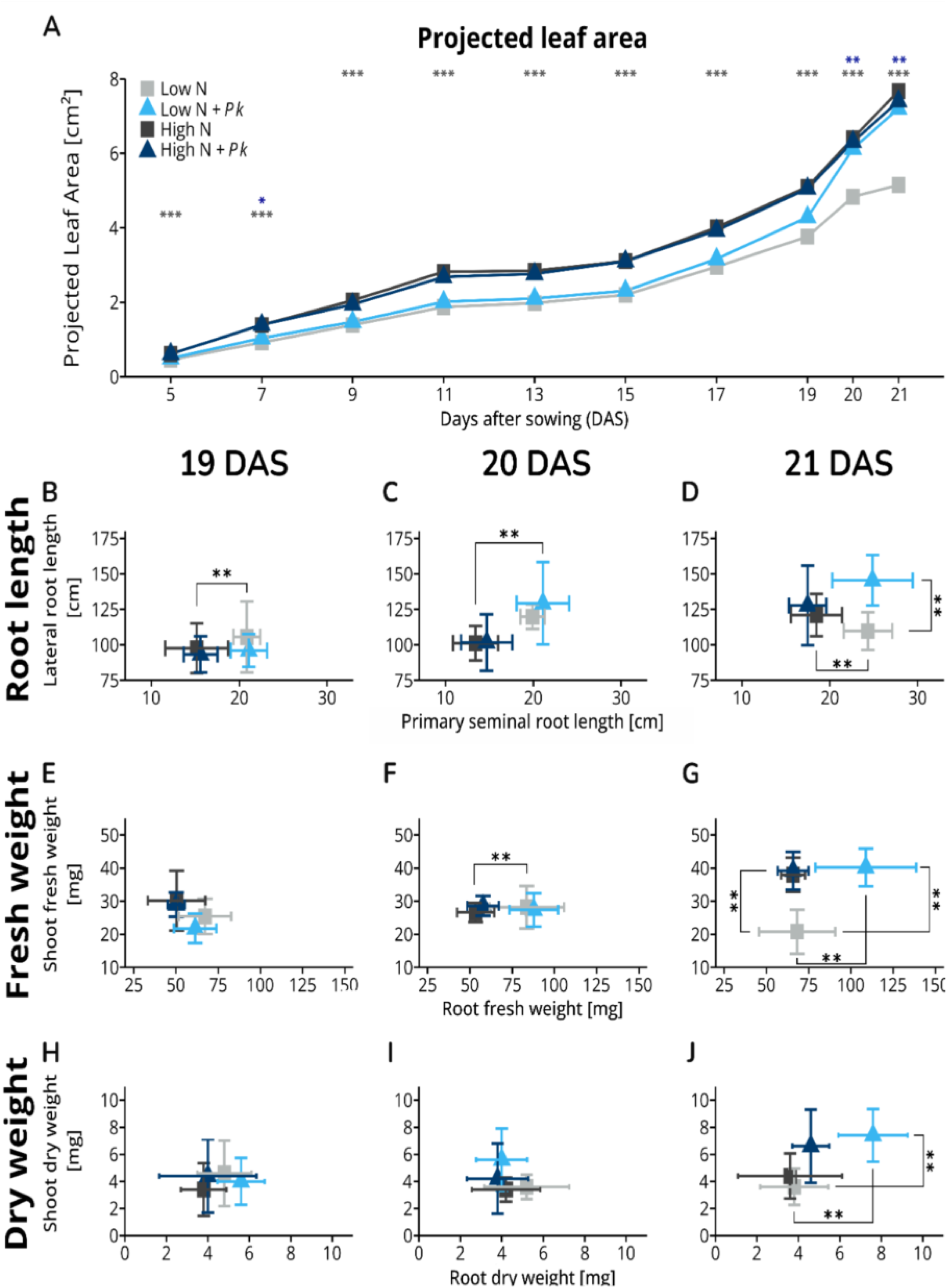
Phenotype overview 21 DAS for harvest I (a comparison of plant growth in all 3 independent experiments is shown in Supplemental Fig S3). A) Non-invasive shoot phenotyping of the time-series. (B -D) Root length distribution with Primary seminal root length (x axis) lateral root length (y axis); (E - G) Fresh weight plot with shoot DW (x axis) shoot FW (y axis); (H - J) Dry weight plot with root biomass (x axis), shoot biomass (y axis). Coloured asterisks in (A) indicate treatments significantly different from low-N controls. Statistical comparisons were performed using two-way ANOVA followed by Tukey’s honestly significant difference (HSD) test. Data are shown as means, with error bars in plots B–J representing ±SD.

Invasive root system analysis revealed a significant increase in total root length (TRL) under low-N + Pk at 21 DAS, with no changes noted at 19 and 20 DAS. Dissecting the root system in primary seminal roots (PSR) and lateral roots (LR), we found that PSR lengths respond to nitrogen availability but remain unaffected by *Pk* inoculation at all time points (Fig. 1 b – d). Notably, low-N + *Pk* plants LR length significantly increased at 21 DAS (Fig. 1 d). These results suggest that *Pk* inoculation induces root architectural changes, potentially facilitating increased surface area for interaction with beneficial microbes and nutrient uptake.

The fresh weight (FW) response to low-N was first visible in the root than the shoot. At 20 DAS, plants under low-N showed a significant increase in root FW compared to those under high-Nitrogen conditions (Fig. 1 f), whereas this trend shifted to a significant increase in shoot FW at 21 DAS (Fig. 1 g). Thus, *Pk* inoculation resulted in a significant FW increase in both root and shoot by 59.5% and 93.3%, respectively, at 21 DAS compared to their uninoculated controls (Fig. 1 f).

Dry weight (DW) did not reveal noticeable differences between the two N conditions (Fig. 1 h, i). However, *Pk* inoculation under low-Nitrogen conditions led to a significant increase in DW in both roots and shoots by 100% compared to their controls at 21 DAS (Fig. 1 j). *Pk* inoculation did not affect plant DW under high-Nitrogen conditions at any time point.

Considering the results in figure 1, the increase in biomass (DW) after *Pk* inoculation under low nitrogen conditions corresponds with the changes observed in FW (Fig. 1 g). Furthermore, the significant increase in root DW aligns with the significant increase in LR length (Fig. 1 d) and, for shoots, with the PLA (Fig. 1 a). Hence, we propose that *Pk* inoculation only operates within a specific range of nitrogen concentrations, enhancing overall plant development in nitrogen-deficient environments.

### Increased N content in Brachypodium after inoculation with *Pk* in a low-N environment

The relative N content (N per milligram (mg) DW) of high-N control plants exhibited significant increases in both roots and shoots compared to low-N control plants (Fig. 2 a – c). While inoculation with *Pk* under low nitrogen conditions did not affect the relative plant N content at 19 and 20 DAS (Fig. 2 a, b), a significant increase in relative shoot N content was observed at 21 DAS (Fig. 2 c). Inoculation with *Pk* under high nitrogen conditions did not impact relative plant N content. The absolute plant N content did not differ between low and high-N control plants at all time points (Fig. 2 d – f). Brachypodium inoculated with *Pk* under low-N conditions led to a significant increase in absolute N content in roots (Fig. 2 f), similarly shoot absolute N content also increased and was comparable to that of plants grown under high-N.

**Fig. 2.**
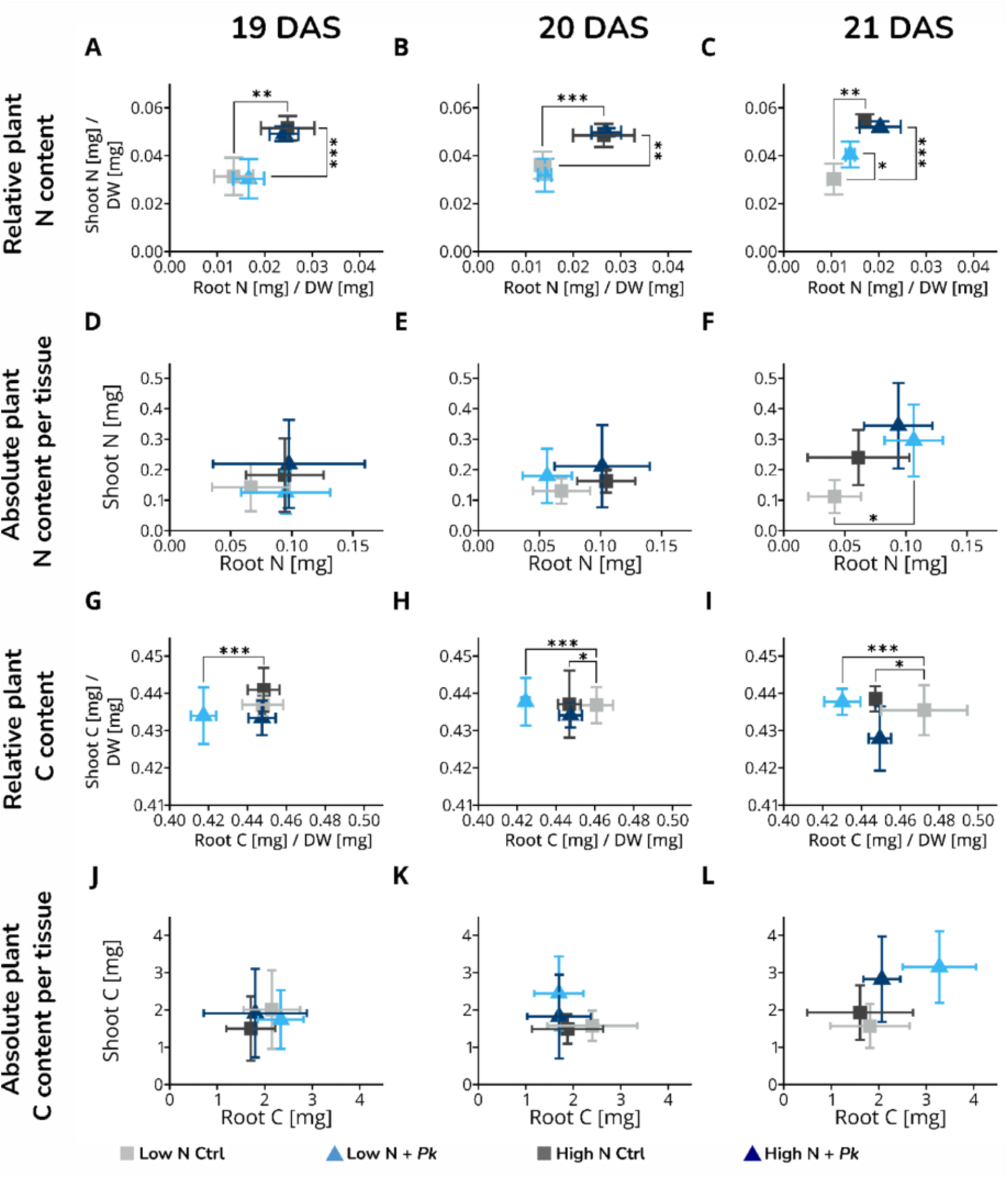
Time-resolved measurements of nitrogen and carbon content in plants at 19, 20, and 21 days after sowing (DAS), shown from left to right across columns. (A–C) Relative nitrogen content (per mg dry weight); (D–F) absolute nitrogen content (mg per tissue); (G–I) relative carbon content (per mg dry weight); (J–L) absolute carbon content (mg per tissue). Statistical comparisons were performed using two-way ANOVA followed by Tukey’s honestly significant difference (HSD) test. Data are shown as means with error bars ±SD.

The relative carbon (C) content (C per milligram (mg) DW) of low and high-N control plants showed no differences in both roots and shoots at 19 DAS (Fig. 2 g). Low-N control plants significantly increased the relative root C content over time compared to high-N control plants (Fig. 2 h, i). The relative shoot C content did not respond to either nitrogen concentration or *Pk* inoculation at all time points (Fig. 2 g – i). Notably, the relative root C content was significantly reduced after inoculation of Brachypodium with *Pk* under low nitrogen compared to both low and high-N plants at all time points (Fig. 2 g – i). The absolute plant C content was not significantly altered at any time point (Fig. 2 j – l).

These results suggest that *Pk* inoculation under low-N conditions increased the overall N content in plants, with a significant increase in relative N content in shoots allowing improved shoot development in a N-deficient environment. Together with the observed increase in absolute root N content after inoculation of Brachypodium with *Pk* under low-N, we hypothesized that *Pk* might be a nitrogen-fixing organism. Significant decrease in the relative C content in the root suggest that there might be ‘communication’ between Brachypodium roots and *Pk* via root exudates, either attracting the beneficial Pseudomonas strain or in exchange for the provided N.

### *Pseudomonas koreensis* can grow on Nitrogen deficient medium *in vitro* but changes to the ^15^N/^14^N isotopic distribution of inoculated plants were not significant

To investigate the N-fixing hypothesis, we submitted *Pk* to zero N conditions in an *in vitro* plate assay. *Pk* was able to grow on N-free medium, while an *E. coli* control could not. In addition, we saw that *P. syringae* and *P. taiwanensis* could also form colonies on N-free medium (Fig. 3). This supports the hypothesis, that *Pk* can fix atmospheric N at least for its own use.

**Fig. 3.**
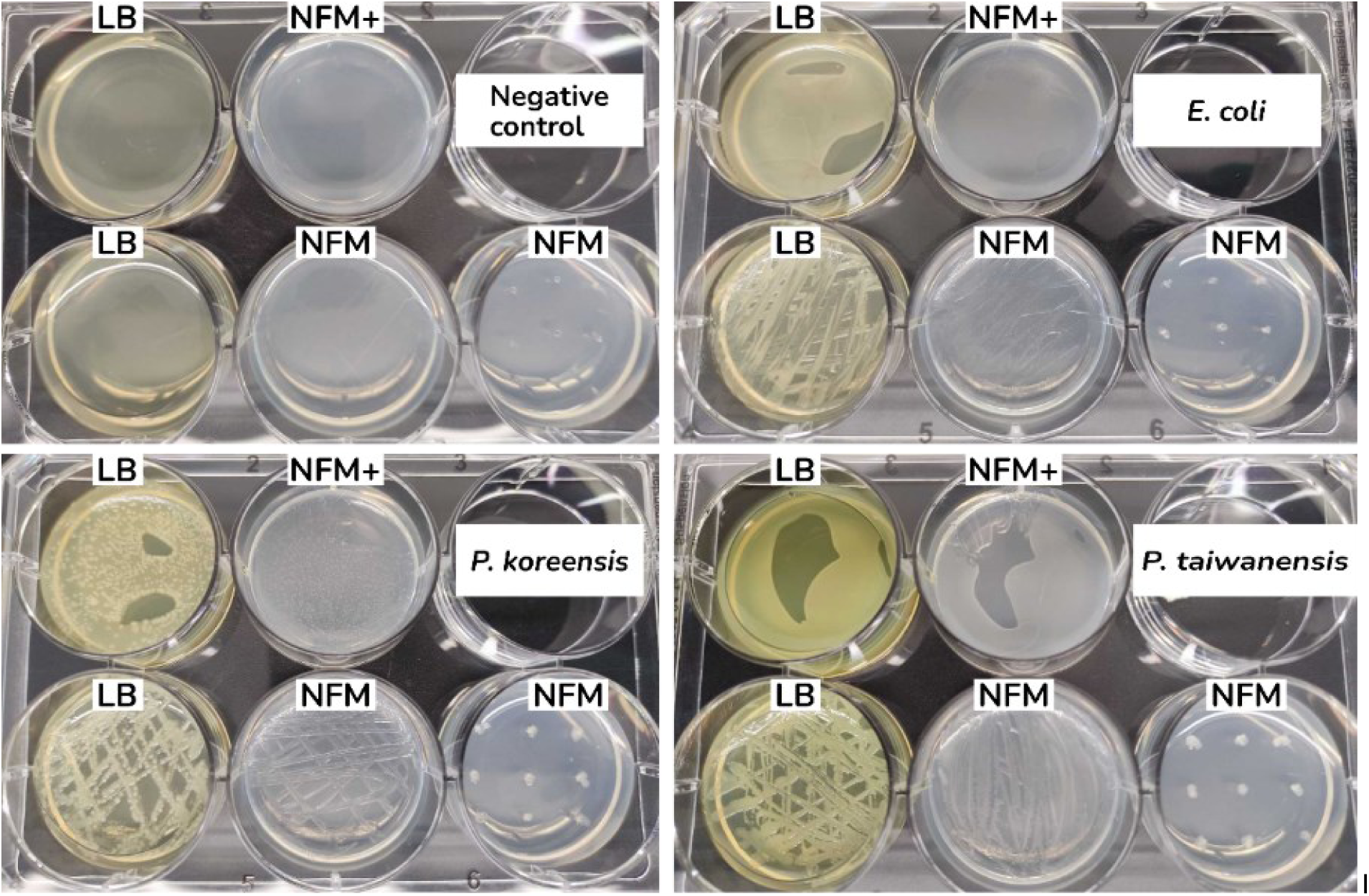
Representative 6-well plates from the nitrogen-fixation plate assay. Wells 1 and 2 contain LB medium; well 3 contains nitrogen-free medium supplemented with N (NFM+); wells 4 and 5 contain nitrogen-free medium (NFM). Well 4 was surface inoculated with bacteria. In well 5, bacteria were introduced by puncturing the agar.

Accordingly, we expected that plants grown together with N-fixing organisms under N-limiting conditions would show a decreased δ^15^N signature vs. air compared to control plants grown without N-fixing organisms (Unkovich, 2013) due to the low δ^15^N signature of biologically fixed N.

Measuring the δ^15^N signature of Brachypodium (Supplemental methods), after inoculation with *Pk,* under three N availability regimes revealed a trend to decreased δ^15^N in the zero N condition in roots (29.1%) and shoots (12.8%) and in low-N conditions in both root (27.3%) and shoot tissues (11.1%) compared to their uninoculated control (Supplemental Fig. 6 a, b). Under sufficient N, only shoots showed a trend of decreased δ^15^N signature (Supplemental Fig. 6 a, b). It seems plausible to assume that under these conditions, N fixation does not occur to any significant extent, and that the response in shoots was due to enhanced biological discrimination against ^15^N. However, the variation among our biological replicates was very large, leading to inconclusive results for *in planta* N -fixation. Studies with a higher gradation of N conditions could potentially provide further insights.

### Molecular adaptations of low-N Brachypodium plants to the presence of *Pk*

To investigate the plant-microbe interaction on molecular level we performed a time resolved investigation of lipidomic changes as lipids are one of the main components of plant cell membranes in addition to many other functions (19, 20 and 21 DAS; Harvest I, Supplemental Fig. 3). At the last time-point we additionally conducted a proteomic analysis as a complementary dataset to understand the plant adaptation to the presence of *Pk* (Harvest II, Supplemental Fig. 3).

### The lipidome of Brachypodium mainly responds to different N levels, with very few *Pk*-induced lipid changes

Principal component analysis (PCA) of all lipids showed differences between high-N controls and low-N controls at all timepoints. On a general level, inoculation with *Pk* did not seem to affect the lipidome under any N condition (Fig. 4 a, c, e).

**Fig. 4.**
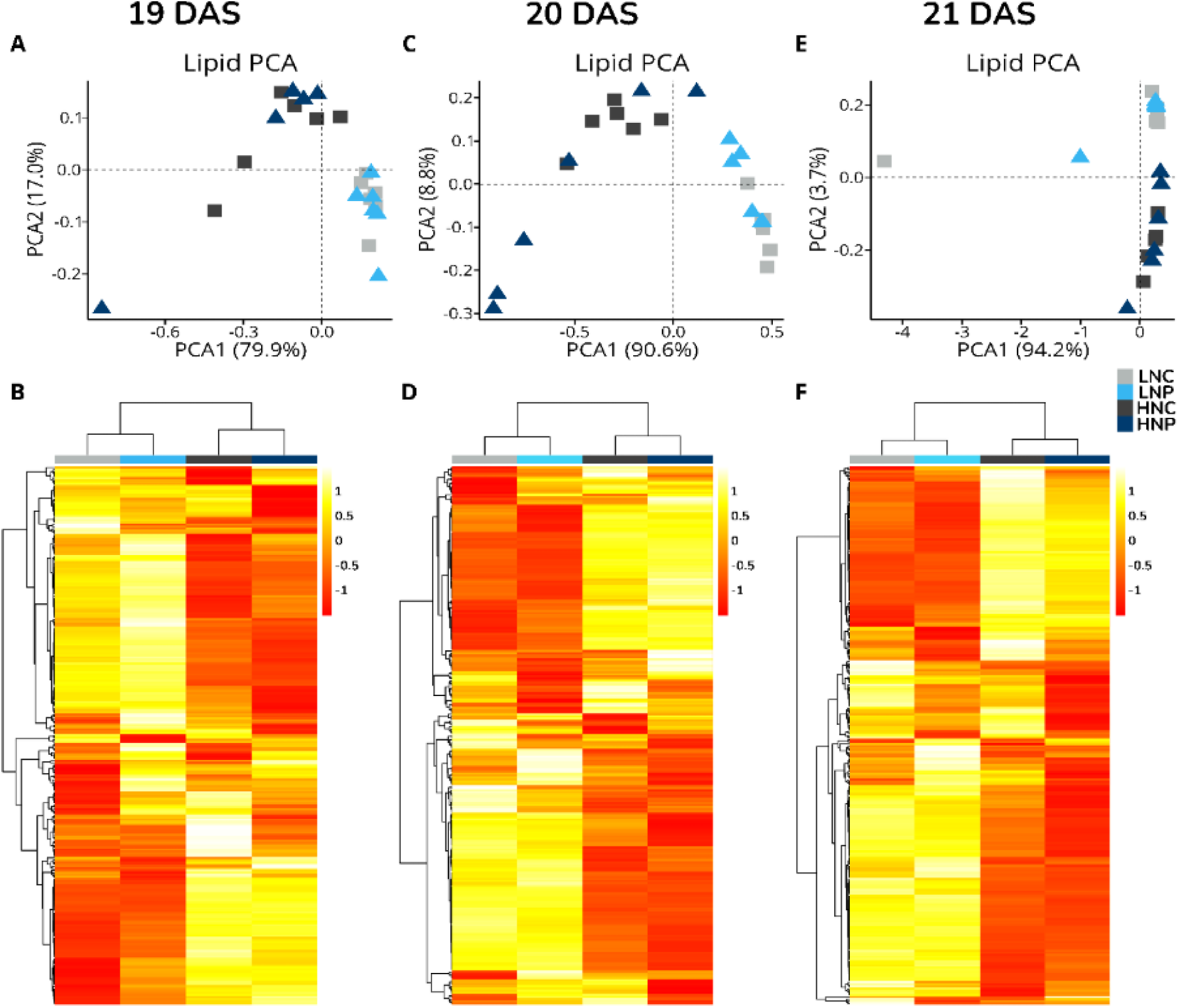
PCA and Heatmaps of root Lipids 19, 20 and 21 DAS (time-resolved). All manually curated lipids (255 features) are mapped using hierarchical cluster analysis (HCA), Pearson distance measurement and average distance. Abundances were log10 transformed and scaled using Pareto scaling. Heatmaps were created using metaboanalyst.com (www.metaboanalyst.com; Xia et al., 2009).

A closer look at the heatmaps (Fig. 4 b, d, f) show that inoculation with *Pk* seems to mildly affect the abundance of several lipids, but most of these changes are not significant. Only 25 lipids out of 255 identified features, showed significant change in abundance due to *Pk* inoculation. The affected lipid classes are the following: Acyl diacylglycerol (ADGGA) (4), ceramide (Cer) (3), diacylglycerol (DG) (1), digalactosyldiacylglycerol (DGDG) (2), lysophosphatidylcholine (LPC) (1), monogalactosyldiacylglycerol (MGDG) (3), phosphatidylcholine (PC) (2), phosphatidylethanolamine (PE) (4), phosphatidylinositol (PI) (1), sulfoquinovosyl diacylglycerol (SQDG) (1) and triacylglycerol (TG) (3).

At 19 DAS, we identified 19 lipids with significantly different abundances between inoculated and non-inoculated samples under low-N, representing the strongest response across all time points. In contrast, only 4 lipids had significantly different abundance at 20 and 21 DAS. These results suggest that major changes in the lipid profile may occur before 19 DAS and therefore warrant further investigation. The time-resolved lipid responses can be found in the supplemental figure 7, and the implications are discussed below.

### Brachypodium proteome is influenced after *Pk* inoculation under limited N

The isolated proteins were analyzed for two purposes: (i) to confirm the inoculation process by searching for bacterial proteins against the Pk database as a qualitative control, and (ii) to investigate how Brachypodium protein abundances were altered by Pk inoculation under the two nitrogen conditions.*Pk* specific proteins confirm the inoculation procedure

A qualitative analysis using the *Pseudomonas koreensis* non-redundant protein library was used to identify *Pk* proteins in our extracts and to determine their presence across samples (Harvest II, Supplemental Fig 3). In inoculated samples, we detected 30 *Pk*-specific proteins and 4 proteins with sequences conserved across multiple bacterial species (Supplemental Fig. 8). In contrast, non-inoculated samples contained no *Pk* specific proteins in most cases. Four control samples contained a small number of *Pk* specific proteins (#2, #4, #10, #14), but markedly less than the inoculated samples (Supplemental Figure 8).

### General overview of the Brachypodium proteome

A total of 5660 and 6656 unique proteins were identified in the root and shoot proteomes, respectively. Of these, 3137 (roots) and 37898 proteins (shoots) were consistently quantified across all samples (Fig. 5 a, b). The number of differentially abundant proteins is listed in Supplemental table S2.

**Fig. 5.**
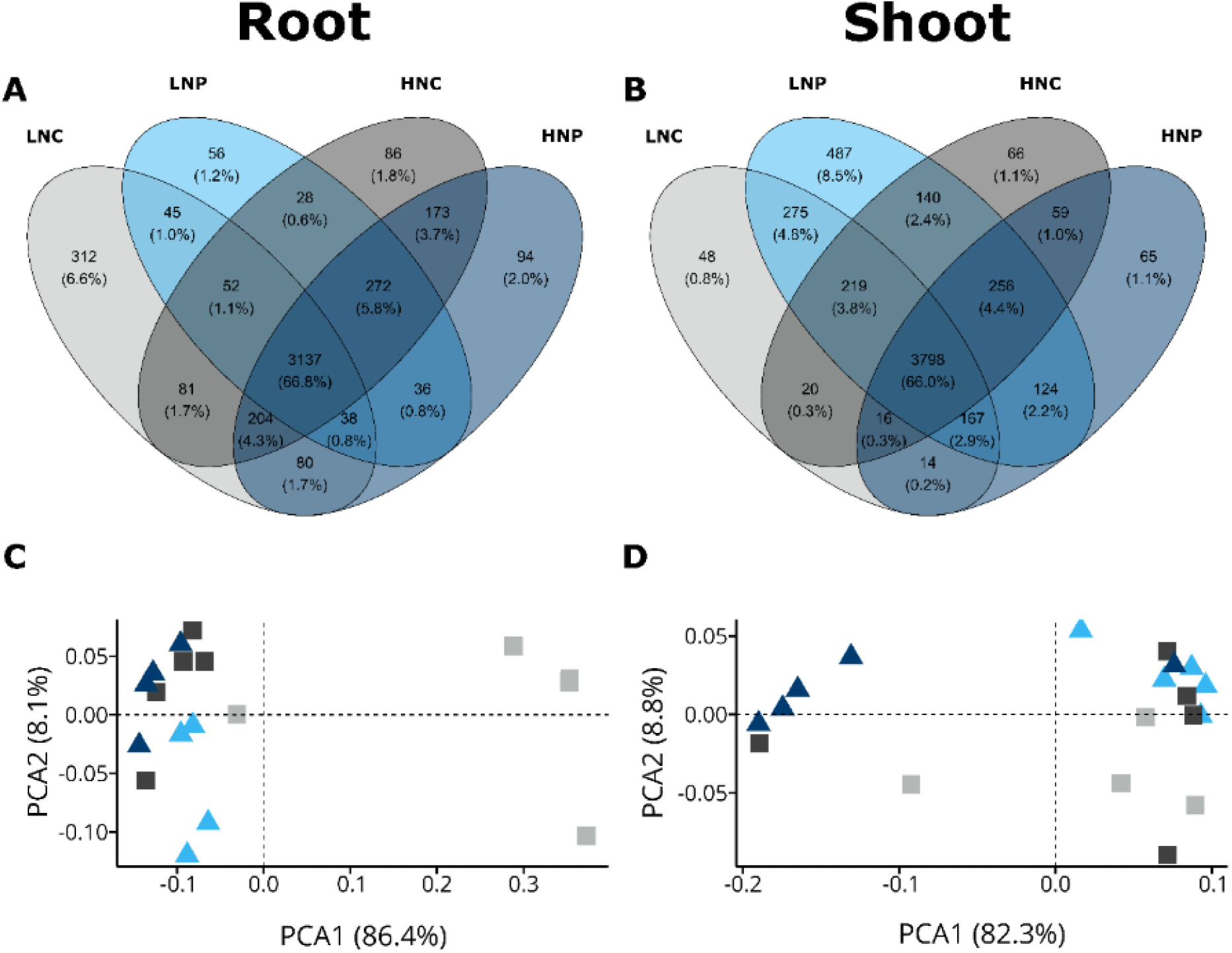
Proteomics data overview. (A, B) Venn diagrams of all proteins with 100% valid values in at least one condition detected in roots (A) and shoots (B). (C, D) Principal component analysis (PCA) of the whole-proteome datasets for roots (C) and shoots (D), based on proteins with 100% valid values in all conditions.

Thus, we conclude that three weeks post inoculation N availability remains the major driver of differential protein expression, although N-limitation enhances the response to *Pk*.

The proteins with 100% valid values (vv) identified in root and shoot tissues are visualized in the principal component analysis (Fig. 5 c).

The volcano plots in Supplemental Fig. S9, show that inoculation under low-N results in substantially more significant protein changes than under high-N. With this in mind, we will mainly focus on the low-N comparison of inoculated vs non inoculated plants in the following figures.

We next compared the biological processes involved in the high-N versus low-N response with those affected by inoculation at low-N (Supplemental Fig. S10). We found that there are more pathways involved in the root response (23) than in shoot (7). In addition, the N response (high vs low) involved different biological processes than the response to the presence of *Pk* (from the top four processes: monoatomic ion transmembrane transport (only lowN+*Pk* vs lowN), cytoplasmic translation (both comparisons), nitrogen compound transport (only High-N vs Low-N), ribonuclein complex biogenesis (both comparisons); Supplemental figure S11).

To obtain a more detailed picture of the systemic response to the presence of *Pk* under low-N, we investigated protein groups which function in processes that we hypothesized to be involved in the response to *Pk* and N. Specifically, we focused on enzymes involved in the lipid pathway (to complement the lipidomic results), kinases as indicators of signalling processes involved in adaptation to both to N and *Pk* as well as N-related metabolic proteins and transporters.

### Few marked proteins involved in lipid metabolism respond to the presence of *Pk*

Following up on the lipidomic results, we investigated whether proteins involved in lipid metabolism (synthesis and degradation) respond to the presence of *Pk*.

Adjustment in lipid-related proteins in the acetyl-CoA synthesis and acetyl-CoA carboxylation pathways, as well as the mitochondrial and plastidial fatty acid synthesis pathways (mtFAS, ptFAS), in the Low-N + *Pk* vs Low-N comparison closely resemble the ratio of High-N vs Low-N control (Fig. 6).

**Fig. 6.**
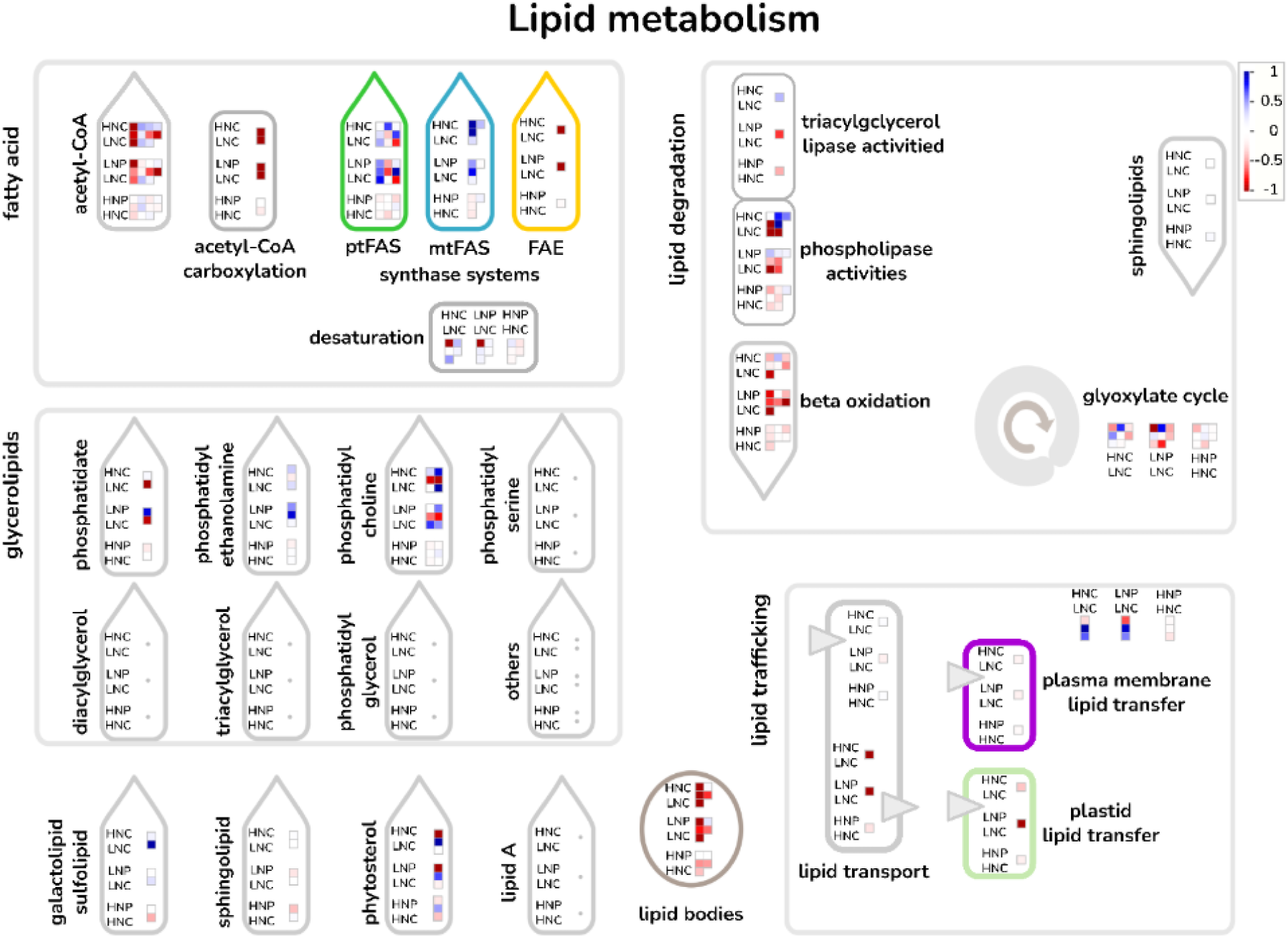
Ratios of proteins involved in lipid metabolism in roots, depicted as heatmap with blue indicating higher abundance and red indicating lower abundance. Data shown represent 100% valid values (vv). Mapped proteins, log_2_FC and significances (indicated as bold values in the log2FC column) are shown in Supplemental Table 2. Pathway mapping was performed using MapMan version 3.5.1R2 (Thimm et al., 2004). The following pathways were used and modified: X4.5 Lipid metabolism R5.0. Mapping file was created using Mercator4 V2.0 (Schwacke et al., 2019). Abbreviations: ptFAS, plastidial fatty acid synthase; mtFAS, mitochondrial fatty acid synthase; FAE, fatty acid elongase. Individual AGI codes and numerical ratios are listed in supplemental table S1. Comparisons are mentioned on next to each box: HNC / LNC = High-N Control vs. Low-N Control; LNP / LNC = Low-N + *Pk* vs. Low-N Control; HNP / HNC = High-N + *Pk* vs. High-N Control.

In context of the lipid degradation pathways, *Pk* inoculation under low-N downregulated the abundance of triacylglycerol lipase (I1HLP4_BRADI), in contrast to the upregulation observed in the High-N versus Low-N comparison.

Similarly, the non-specific phospholipase C (nPLC) (A0A0Q3GLT1_BRADI) involved in phospholipid degradation showed reduced abundance compared to its expression at low-N control.

In conclusion, Pk primarily influences lipid degradation under low-N conditions.

### Kinase families in Brachypodium respond to N availability and *Pk* inoculation

We compared the abundance of the various kinase families (Supplemental Fig. 12, Supplemental Table 1), including many that have an extracellular domain and function across the plasma membrane. Here, we highlight the Ca²⁺/calmodulin-dependent protein kinase (CAMK) group, with particular focus on the SnRK1 kinase complex, a regulatory hotspot for energy homeostasis and stress responses including nutrient signaling (Broeckx et al., 2016; Crepin and Rolland, 2019).

In roots, the SnRK2 SNF1-related protein kinase SAPK6 (I1IAE4_BRADI) was more abundant in high-N Controls compared to low-N Controls (1.05 log2FC), which was also observed after inoculation of low-N plants with *Pk* (0.54 log2FC) and mildly after inoculation of high-N plants (0.29 log2FC). The regulatory subunit beta of SNF1-related SnRK1 kinase complex (I1HI64_BRADI) accumulated under high-N controls vs. low-N controls (0.83 log2FC), whereas SAPK3 (I1I692_BRADI), SAPK7 (I1IXT7_BRADI) and SAPK8 (I1GMZ0_BRADI) were generally reduced compared to low-N controls, with only slight reductions in high-N + *Pk.* In contrast, the regulatory subunit betagamma of SNF1-related SnRK1 kinase complex (I1GKM6_BRADI) decreased strongly under high-N controls compared to low-N controls (-0.73 log2FC) and after inoculation of low-N plants (-0.91 log2FC). However, the regulatory subunit gamma of SNF1-related SNRK1 kinase complex (I1HNY1_BRADI), remained unchanged in high-N controls compared to low-N controls (0.02 log2FC), but increased moderately in low-N inoculated plants compared to low-N control (0.56 log2FC). It was also moderately less abundant after inoculation of high-N plants (-0.43 log2FC) compared to high-N controls.

The biggest differences were observed in CDPK protein kinases. Expression of two CDPKs (I1GMG8_BRADI; I1H269_BRADI), decreased under high-N controls vs. low-N controls (-0.39; -0.18 log2FC, respectively) but increased after low-N + *Pk* inoculation (0.25; 0.37 log₂FC), with little change under high-N + *Pk*. Another CDPK (I1J0Z2_BRADI) and the CRK kinase (I1GX02_BRADI) were strongly reduced under high-N versus low-N (–1.16; –0.9 log₂FC) and low-N + Pk (–0.75; –1.11 log₂FC), but remained unchanged under high-N + Pk.

As part of the CMGC kinase superfamily, the MAP kinase I1I5N8_BRADI was found to be moderately less abundant in the high-N Controls compared to low-N controls (-0.51 log2FC), much less abundant after inoculation of low-N plants (-1.33 log2FC) and unchanged after inoculation of high-N plants.

Overall, in roots the abundance ratios of many kinases under high-N versus low-N were mirrored in low-N + Pk versus low-N, suggesting overlapping regulation. Exceptions from this trend are considered potential candidates for adaptation to beneficial microbes.

In the shoot, on the other hand first we see fewer significantly regulated kinases and second more of them have opposite behaviour between the high-N control vs low-N control, and low-N + *Pk* vs low-N control. This is in line with our hypothesis that they might be involved in the response to PGPB.

In the case of L-type lectin receptor kinases which are implicated in plant immunity (Wang & Bouwmeester, 2017), we see higher abundance in high-N than in low-N, but at low-N + *Pk* decreases the abundance of kinase A0A0Q3KPD2_BRADI.

An opposite behaviour was seen in a representative of the CMGC kinase superfamily (comprising of CDK, MAPK, GSK3 and CLK families), I1I8I8_BRADI, the CAMK kinase superfamily proteins (I1H626_BRADI, I1HPI9_BRADI and I1GKM6_BRADI) and the LRR kinases (I1ID77_BRADI, A0A0Q3EYZ3_BRADI and A0A2K2D9S6_BRADI). These kinases are downregulated under high nitrogen compared to deficiency, but upregulated at low-N in the presence of *Pk*. We speculate that these kinases (and other regulated in a similar manner) represent internal plant regulatory adaptations to presence of the beneficial microbe.

### *Pk* inoculated *Brachypodium* decreases N-deficiency symptoms in low-N roots but increases N assimilation enzymes in the shoot

Guided by the increase of N in the plants (Fig. 2 c, f) we checked amendments to the abundance of proteins involved in N uptake and central N metabolism. Transporters and enzymes of the central N metabolism of plants were of particular interest, as they are the main actors in assimilating inorganic nitrogen into organic molecules in plants. The proteome dataset allowed us to confirm that (i) the low-N plants do indeed show the text-book N deficiency symptoms, as well as (ii) inoculation altered the abundance of N metabolism proteins under both N conditions. For visualization, we use the low-N control of root and shoots as the baseline for comparison for the respective tissue.

The different N-conditions showed the expected change in protein abundance. In roots, high-N control (HNC) plants showed lower abundance of NRT and AMT transporters compared to low-N control (LNC) plants (Fig. 7, roots). An exception was one NRT transporter (I1H1D1_BRADI), which showed higher abundance in HNC roots.

**Fig. 7.**
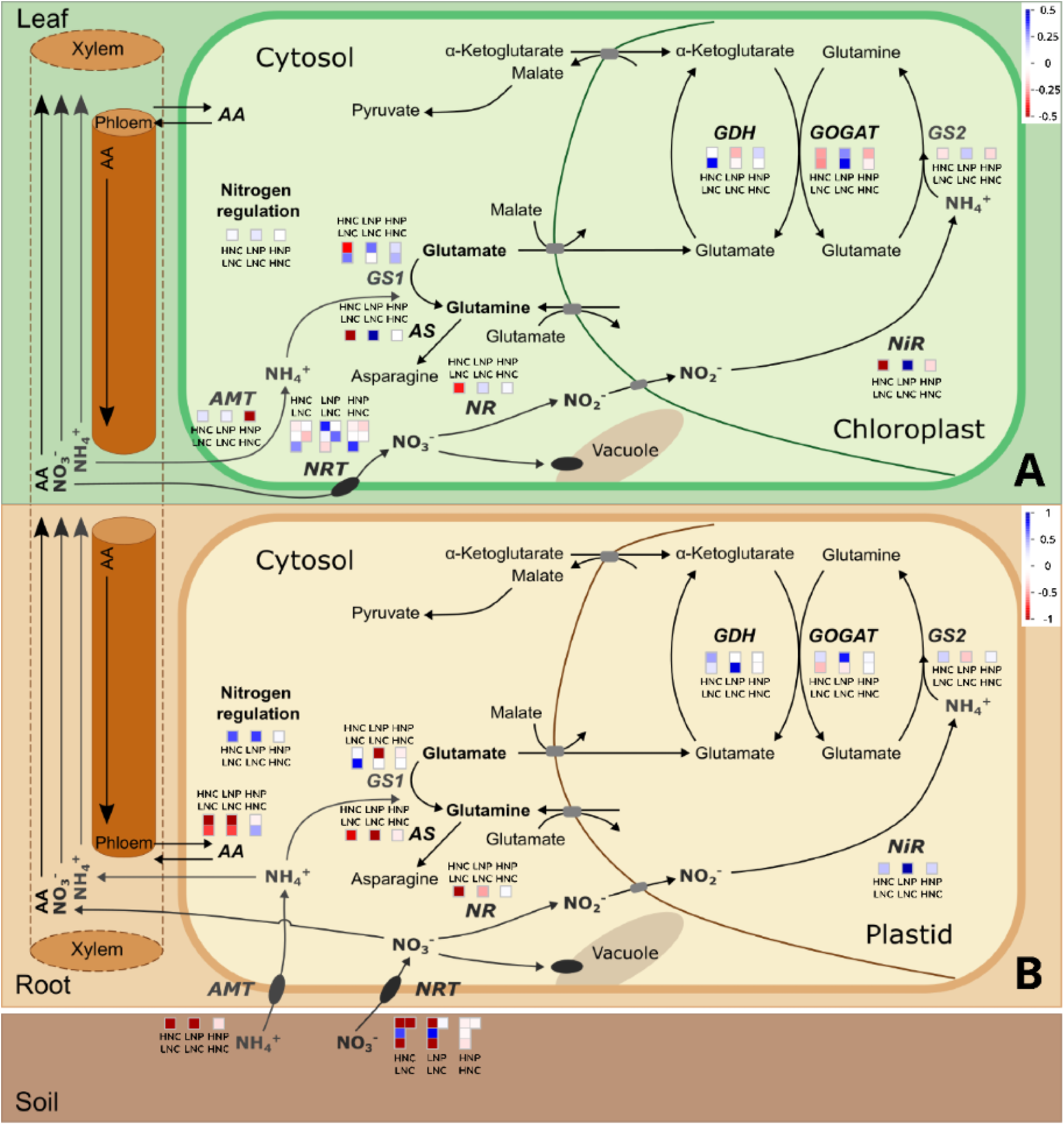
Proteins of Central N metabolism in roots and shoots of the comparison low-N + *Pk* vs. low-N Control, depicted as heatmap with blue indicating higher abundance and red indicating lower abundance. Data represented are 100% valid values. Mapped proteins, log2FC and significances (indicated with asterisks in log2FC column) are shown in supplemental table 2. MapMan version 3.5.1R2 (Thimm et al., 2004) was used. The following pathways were used and modified from Feng et al., 2020: X4.1 N-uptake R1.0. Modified pathways will be submitted to the MapMan store (https://mapman.gabipd.org/mapmanstore). Mapping file was created using Mercator4 V2.0 (Schwacke et al., 2019). Comparisons are mentioned on top or below of each box: HNC / LNC = High-N Control vs. Low-N Control; LNP / LNC = Low-N + *Pk* vs. Low-N Control; HNP / HNC = High-N + *Pk* vs. High-N Control. Abbreviations: AA, Amino acid; AMT, ammonium transporter family; AS, Asparagine synthase; GDH, Glutamate dehydrogenase; GOGAT, glutamine oxoglutarate aminotransferase; GS1, (cytosolic) Glutamine synthetase 1 isoforms; GS2, (plastidial) Glutamine synthetase 2 isoforms; NiR, Nitrite reductase; NR, Nitrate reductase; NRT, Nitrate transporter family.

We identified both Fd- and NADH-GOGAT isoforms in the roots, which was surprising because plants were grown in sand and thus the roots were protected from light (Yang et al., 2016). Fd-GOGAT abundance was slightly higher in HNC roots, whereas abundance of NADH-GOGAT was lower. The Nitrogen regulatory protein P-II was also more abundant in the high-N control roots, compared to the low-N.

In shoots, N availability similarly influenced proteins of the NO₃⁻ assimilation pathway, including NRT, NR, NiR, GS2, and both GOGAT isoforms (Fig. 7, Supplemental Table 2). Cytosolic glutamine synthases were differentially regulated: the low-affinity isoform GLN1;2 (I1H7X9_BRADI) was less abundant under high-N, whereas the high-affinity isoform GLN1;1 (I1IFI7_BRADI) was more abundant.

In the comparison of low-N + *Pk* vs low-N control plants (Fig. 7) we found lesser and unchanged abundance of 2 NRT transporters in the inoculated roots compared to the low-N control roots (–1.47 and 0.13 log2FC). Uncharacterized protein (I1I6K3_BRADI); MFS domain-containing protein (I1I5R4_BRADI) and NAR2.1 (I1IB70_BRADI) showed no change, while MFS domain-containing protein (I1H1D1_BRADI) was higher abundant in low-N + *Pk* roots. The nitrate reductase (NR. A0A0Q3FHC7_BRADI) was less abundant, but the nitrite reductase (NiR, I1IEV4_BRADI) was more abundant in low-N inoculated roots vs the low-N control. The plastidial GS2 (I1H7X9_BRADI) is less abundant, but there is higher abundance of the Fd-GOGAT isoform (I1GRK9_BRADI) and lower abundance of the NADH-GOGAT isoform (I1HQF1_BRADI) and related enzymes (Fig. 7, root). The detected ammonium transporters (AMT) and several enzymes involved in assimilation of NH_4_^+^ in amino acids are less abundant in the low-N +*Pk* roots. There is strongly increased abundance in the Nitrogen regulatory protein P-II (A0A0Q3MUB2_BRADI) (Fig. 7, root).

In the low-N + *Pk* shoots compared to the low-N control shoots we see consistently increased abundance of the NO_3_^-^ assimilation pathway, as well as increased abundance of enzymes of the NH_4_^+^ assimilation pathway. Enzymes like AS (I1H6K4_BRADI) and GS (I1H7X9_BRADI) involved in the synthesis of amino acids are more abundant (Fig. 7, shoot). This points to increased investment, of protein abundance and perhaps energy, for shoot N uptake and N relocation form the root.

In contrast, in the comparison of high-N roots + *Pk* vs high-N control roots show very mild differences in abundance in the central N metabolism in the roots (Fig. 7, root). Particularly interesting, the nitrogen regulatory P II protein is unchanged. In the shoot there is a higher abundance of a single NRT transporter (I1H1D1_BRADI) and mildly increased abundance of the two detected cytosolic GS isoforms. This mimics the very mild response to *Pk* inoculation already seen in the root.

## Discussion

Our findings demonstrate that *Pk* strongly promotes growth under low-N conditions, with inoculated plants producing nearly twice the biomass of uninoculated controls within 21 days This growth promotion is also reflected by an overall increase in N content in the shoots of low-N + *Pk* plants (whereas the roots show a trend to increased N content that fell below the significance level; Fig. 2 f). These findings suggest either a reallocation of N resources from roots to shoots or an improved root nitrogen use efficiency (NUE), resulting in more biomass per unit N. The shoots of the inoculated plants are bigger but also contain more N per mg dry weight. (Fig. 2 f, d). C content in the shoots showed no difference in the four conditions, whereas the roots of low-N + *Pk* plants showed lowest amount of C.

Accordingly, many proteins in the central N metabolism are adjusted after *Pk* inoculation under low-N, with much weaker effects observed under high-N. Consistently, the proteome PCA mirrored the phenotypic observations: high-N control and high-N + *Pk* roots clustered closely at both molecular (Fig. 5) and phenotypic levels (root length, fresh weight, and C/N content; Fig. 1–2). Notably, the root protein profile of Low-N + *Pk* plants resembled that of high-N controls, indicating that inoculated roots under limiting nitrogen behave more like the high-N plants.

### Nutrients and bacteria influence plant phenotype in different temporal scales

In this study, differences in leaf area due to nitrogen supply were already quantifiable early, at 5 days after sowing (DAS). By contrast, the influence of the plant growth-promoting bacterium Pseudomonas koreensis (*Pk*) became consistently visible only at 19 DAS, i.e., 14 days after inoculation. This is slightly faster than a related study also using Brachypodium at two N conditions, but plants were grown in a gnotobiotic microchamber with liquid medium and inoculated with *Herbaspirillum seropedicae.* In this case, leaf area showed a significant upregulation due to the presence of the bacterium 19 days after inoculation. (Kuang et al 2022).

At the root level, both *Herbaspirillum* and *Pseudomonas* were found to increase root length at low-N, however while the former increased primary root length (Kuang et al, 2022), and the latter increased the root length via lateral roots (Fig. 1 d). This suggests that both the bacterial inoculant and environmental conditions shape the timing of plant responses, but these represent only a fraction of the true complexity of plant-microbe interactions. In natural settings, such interactions are further influenced by dynamic root development (Kawa and Brady, 2022), and shifting microbiome compositions, across root-associated compartments, from the endosphere to the rhizosphere and bulk soil (Singer et al., 2021, Bai et al., 2022). Given the vast diversity of microbes encountered by roots, a wide range of microbe-associated molecular patterns (MAMPs) and damage-associated patterns (DAMPs) are involved in mediating these interactions (Zhou et al., 2020; Ma et al., 2021; Kawa & Brady 2022).

### Multiple mechanisms of *Pk* plant growth promotion and open questions about the plant-microbe interaction

Since the ^15^N data did not unambiguously confirm BNF *in planta*, the increased N content of inoculated plants may also result from enhanced foraging due to the longer root system. At 21 DAS, lateral root length was significantly increased in low-N + *Pk* plants compared to their respective controls. However, in the time-resolved data, lateral root length increases were only evident at 21 DAS, whereas increased N content was first observed for shoots at 19 DAS and roots at 20 DAS (Fig. 2). The ability of *Pk* to fix nitrogen for its own use, together with the timeline of increased plant N content, indicates that we cannot exclude BNF as plant growth promoting mechanism. However, it is unlikely to be the sole driver of the observed growth promotion, as discussed below.

Similarly, in the present study, inoculated roots under low-N have markedly lower C content compared to controls, whereas Kuang et al (2022) observed no such effect. Whether plant-derived C was utilized to promote the establishment of plant-microbe-interaction (Sasse *et al* 2018), remains an open question for future research.

At the lipid level, very few significant differences were detected at 20 and 21 DAS, while at 19 DAS, Brachypodium roots showed various lipidomic responses to *Pk* under low-N. This pattern suggests that lipid remodeling may occur earlier than our observed time window and could play a role in initiating growth promotion and shaping plant–microbe interactions.

### Remodelling of plant energy metabolism to provide resources for abiotic stress tolerance

Overall, lipid profile changes in Brachypodium likely play a subtle but potentially important role in this plant-microbe interaction.

Lipidomic profiling across 19–21 DAS supports the proteomic evidence of adaptive metabolic reprogramming under nitrogen limitation. A transient accumulation of triacylglycerols (TGs) at 19 DAS in *P. koreensis*-inoculated roots suggests temporary carbon storage to fuel downstream nitrogen assimilation and amino acid biosynthesis at 20 and 21 DAS. Concurrently, the phospholipid profile revealed a decline in phosphatidylcholine (PC) and a relative enrichment in phosphatidylethanolamine (PE), indicating membrane remodelling that may facilitate nutrient transporter activity and vesicular trafficking. Elevated lysophosphatidylcholine (LPC 16:0) across all timepoints reflects enhanced phospholipid turnover and membrane fluidity, potentially supporting signalling and stress adaptation. While plastid-associated lipids galactolipids (MGDG, DGDG, SQDG) remained largely stable, modest increases in DGDG 34:4 and SQDG 34:3 at 21 DAS suggest subtle adjustments in plastidial membrane composition, possibly linked to redox balance or phosphate homeostasis. Minimal changes in diacylglycerol (DG), ceramide, and cardiolipin levels imply that membrane lipid signalling and mitochondrial remodelling are not prominently engaged, aligning with a regulated, non-pathogenic symbiosis. Altogether, the lipidome complements the proteomic data, underscoring a coordinated reallocation of carbon and membrane resources to sustain nitrogen assimilation and cellular resilience under low low-N conditions.

It should be noted that lipid annotation in plants remains challenging (Macabuhay et al., 2022). In our dataset, 255 lipid features were detected, many of which changed over time. Re-examination of this dataset should be performed once the lipid databases become more comprehensive in the context of plant-lipids.

The energy and elemental resources provided from lipid degradation would become available to the plant for adapting the proteome and root system architecture profile for low-N conditions. During low-N plants increase the expression of high-affinity transport system (HATS) to enhance the uptake of NO_3_^-^ and NH_4_^+^ from N-deficient soil. Examples include NRT2.1, NRT2.2 and NRT2.4 (which are all regulated by NAR2.1, Kotur et al., 2012; Laugier et al., 2012) and AMT1;1, AMT1;2 (which are both detected in this study) and AMT1;3 (not detected). This points towards increasing Nitrogen Uptake Efficiency, enhancing their ability to capture and utilize limited nitrogen resources efficiently.

Importantly, NAR2.1 not only regulates most NRT2 transporters (except NRT2.7 in A. thaliana) but is also implicated in lateral root development via effects on auxin transport under low nitrate (Wang et al., 2023). LR formation, most likely a combination of NO_3_^-^ uptake and NO_3_^-^ signalling (Sun et al., 2017), is one of the main findings of this study and is detected after the lipid degradation takes place.

## Concluding remarks

Mineral nitrogen fertilizers elicit rapid plant responses within days, whereas bacterial inoculants act more subtly and over extended periods - weeks to months. Upon colonization, the bacteria interact with the native soil microbiomes and potentially persist through self-propagation, enabling plants to engage diverse metabolic pathways and establish associative relationships that enhance resilience under abiotic stress.

It is important to note that this study was conducted in a simplified, closed system involving a single plant–microbe interaction in the absence of a background microbiome, and thus consequently may not be fully representative of an interaction that would have formed under more natural conditions where plants actively shape their microbiomes.

On the other hand, our simplified system allows for unprecedented level of dissection of the plant and microbial response, through time and under a multitude of conditions, giving clear insight into the plasticity of the plant response, which may be difficult to achieve in real-world conditions.

In conclusion, our temporal dissection of abiotic and biotic interactions at the phenotypic and molecular level highlights the need for: (i) evaluation of plant-microbe dynamics in more complex systems (e.g. soil, field). Recent studies in maize have demonstrated that plant genotypes differ in their ability to recruit beneficial microbes under nutrient limitation, and that the underlying genetic determinants can even be leveraged for breeding (Yu et al., 2021; He et al., 2024: He et al., 2025.) Such “consensual” interactions highlight that integrating plant genotype–microbiome interactions into breeding programs could complement or, in some cases, reduce the need for inoculation strategies. (ii) Careful consideration of interaction timelines in experimental design for future implications, particularly during upscaling for application, and (iii) Continued exploration of plant growth-promoting bacteria as tools to improve plant performance under nitrogen limitation. Most importantly, such approaches could contribute to reducing the reliance on synthetic nitrogen fertilizers in sustainable agriculture.

## Supporting information

Sanow et al_Supplemental files and tables

## Acknowledgements

We thank Ben Crossett and Kang-Yu Peng for performing the lipidomics mass spectrometry measurements at Sydney Mass Spectrometry, a core research facility at the University of Sydney. Special thanks to Thusita Rupasinghe, Allene Macabuhay and Alina Ebert for assistance and advice with the lipidomics mass spectrometry analysis. Thanks to Helena Bochmann for assistance with root phenotyping during Harvest I. Stefan Sanow thanks the Jülich-University of Melbourne Postgraduate Academy (JUMPA) program for the joint PhD scholarship. This work was partially funded by the Deutsche Forschungsgemeinschaft (DFG, German Research Foundation) under Germany’s Excellence Strategy – EXC 2070 – 390732324 (PhenoRob)

## Author contributions

SS, RW, MW, UR, GS, PH, AL, BA designed the research and experiments. SS performed the experiments. ES performed N-fixation assay. MM, HW, AB, JKM performed measurements and/or supported data analysis. All authors contributed to write the final manuscript.

## Data availability

The mass spectrometry proteomics data have been deposited to the PRIDE repository (Perez-Riverol et al., 2025).

The metabolomics data have been deposited to MetaboLights (Yurekten et al., 2024).

Both datasets will be made available after peer-review is completed.

