## Supplementary material for "Systems-level Plant Responses Reveal *Pseudomonas*-Mediated Growth Promotion in *Brachypodium* Under Nitrogen Limitation": Sanow et al_Supplemental files and tables

### Supplemental figures

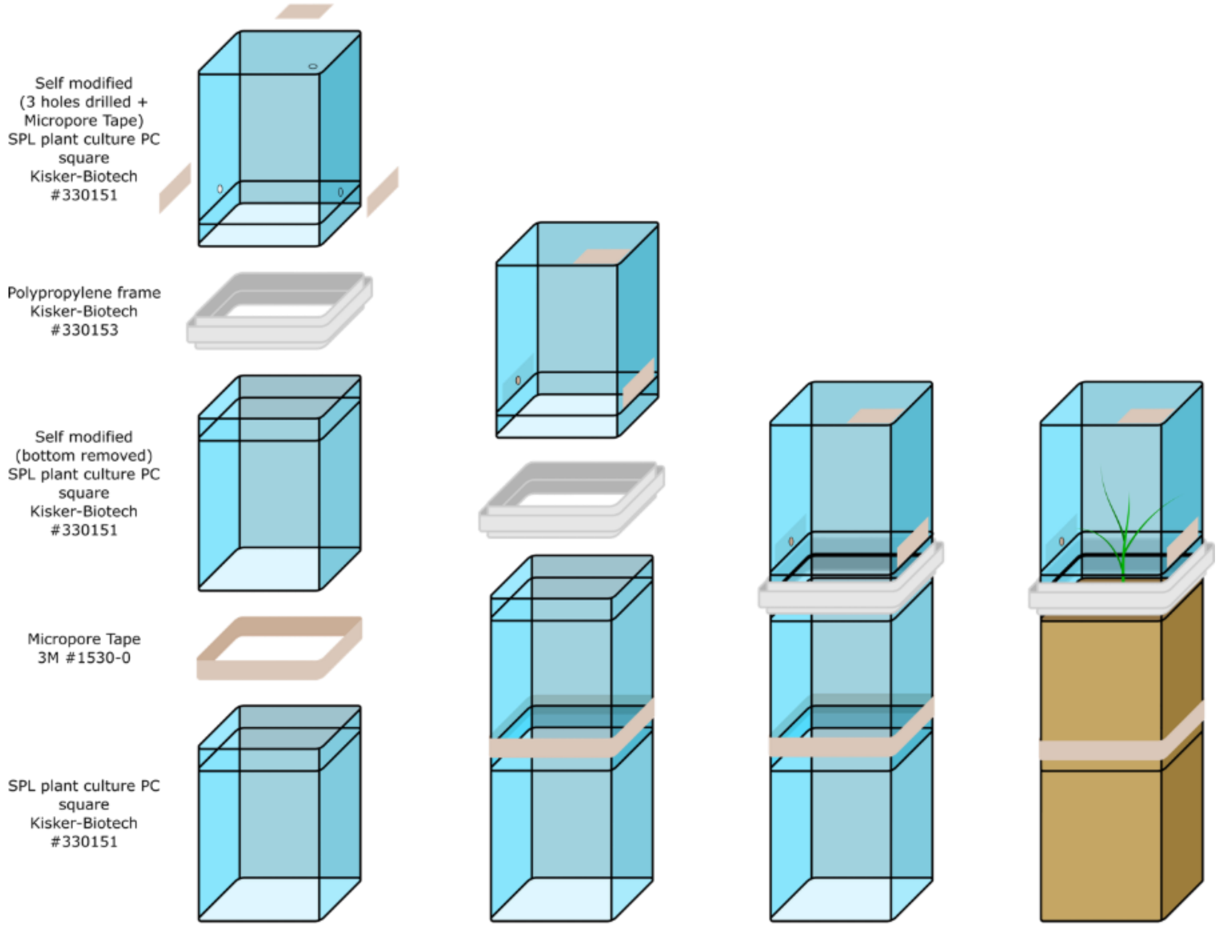


**Supplemental Figure S1**

PCV modification and assembly. Three individual Plant Cultivation Vessels (PCV) were stacked on top of each other to obtain a single, closed system. The middle PCV had its bottom sawed off and was stacked on top of the bottommost PCV, using micropore tape (3M) to fix the two containers. The inverted PCV used as a lid was linked to the middle container using the commercially available connector (#330153, Kisker Biotech). For better gas exchange, a total of 3 holes were drilled in the lid.

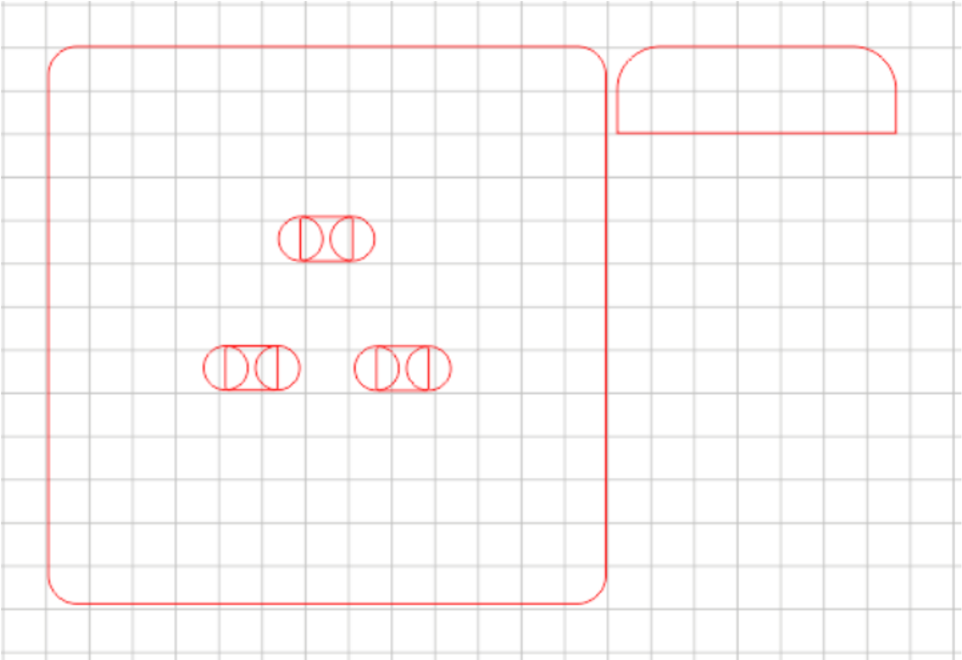

**Supplemental Figure S2**

Sowing template for laser cutter. Left: Base of the sowing template, including 3 holes for fixed seed sowing positions, right: handle, which can be glued to the base for easier operation. The plan for the sowing template, produced by a 3D laser cutter (Trotec, Speedy 100). The sowing template (3 mm thickness) is pressed into the PCV until it is flush with the rim. Then, three holes were poked in the sand using a glass stirring rod, with a stopper at depth of 1.3 cm.

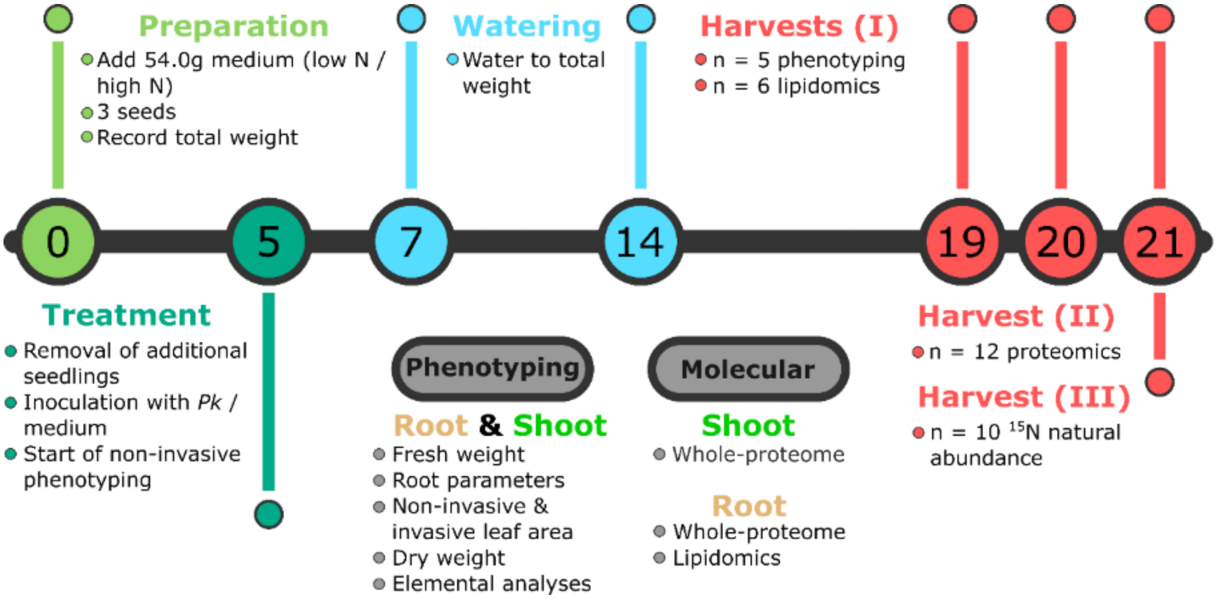


**Supplemental Figure S3**

Schematic overview of the experimental setup. Numbers in the circles represent days after sowing (DAS). Non-invasive phenotyping was carried out every 2 days after 5 DAS, and on each harvest day. Plants were grown under four different conditions: Low-N, Low-N + *Pk*, High-N, High-N + *Pk*. Three independent experiments were harvested: Harvest (I) Time-series includes consecutive harvests on 19, 20, 21 DAS, with n = 5 plants for phenotyping and n = 6 plants for lipidomics per condition and per timepoint; Harvest (II) at 21 DAS with n = 12 plants for proteomics, and Harvest (III) at 21 DAS with n = 5 plants for $^{15}N/^{14}N$ isotope distribution . Abbreviations: PCV = Plant Cultivation Vessel, *Pk* = *Pseudomonas koreensis.*

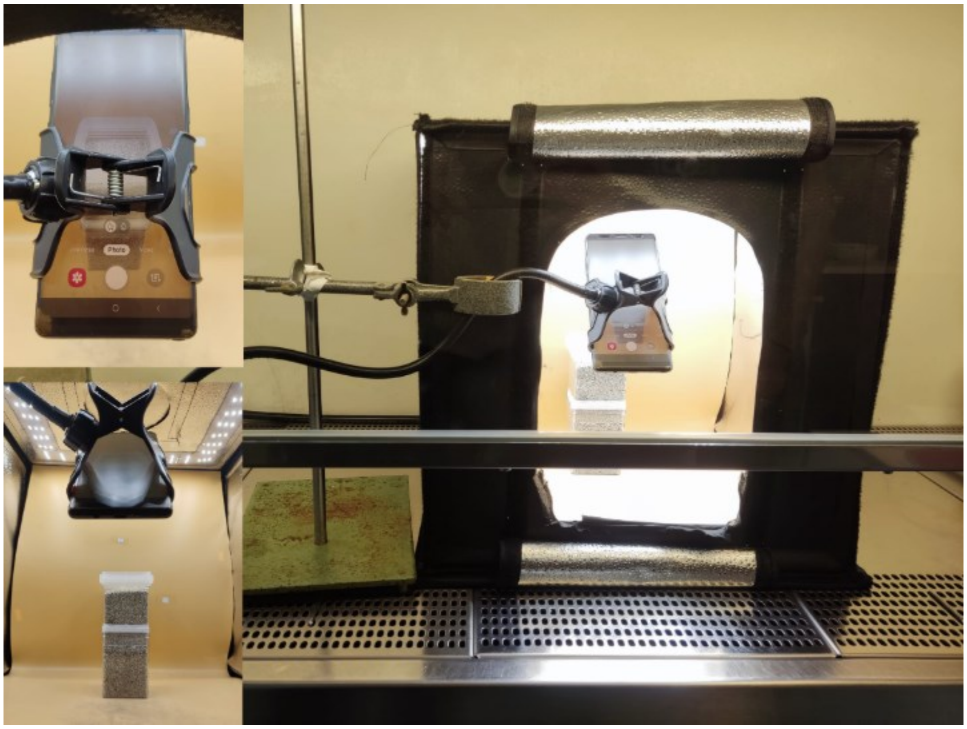

**Supplemental Figure S4**

Non-invasive shoot phenotyping setup inside a biosafety cabinet.

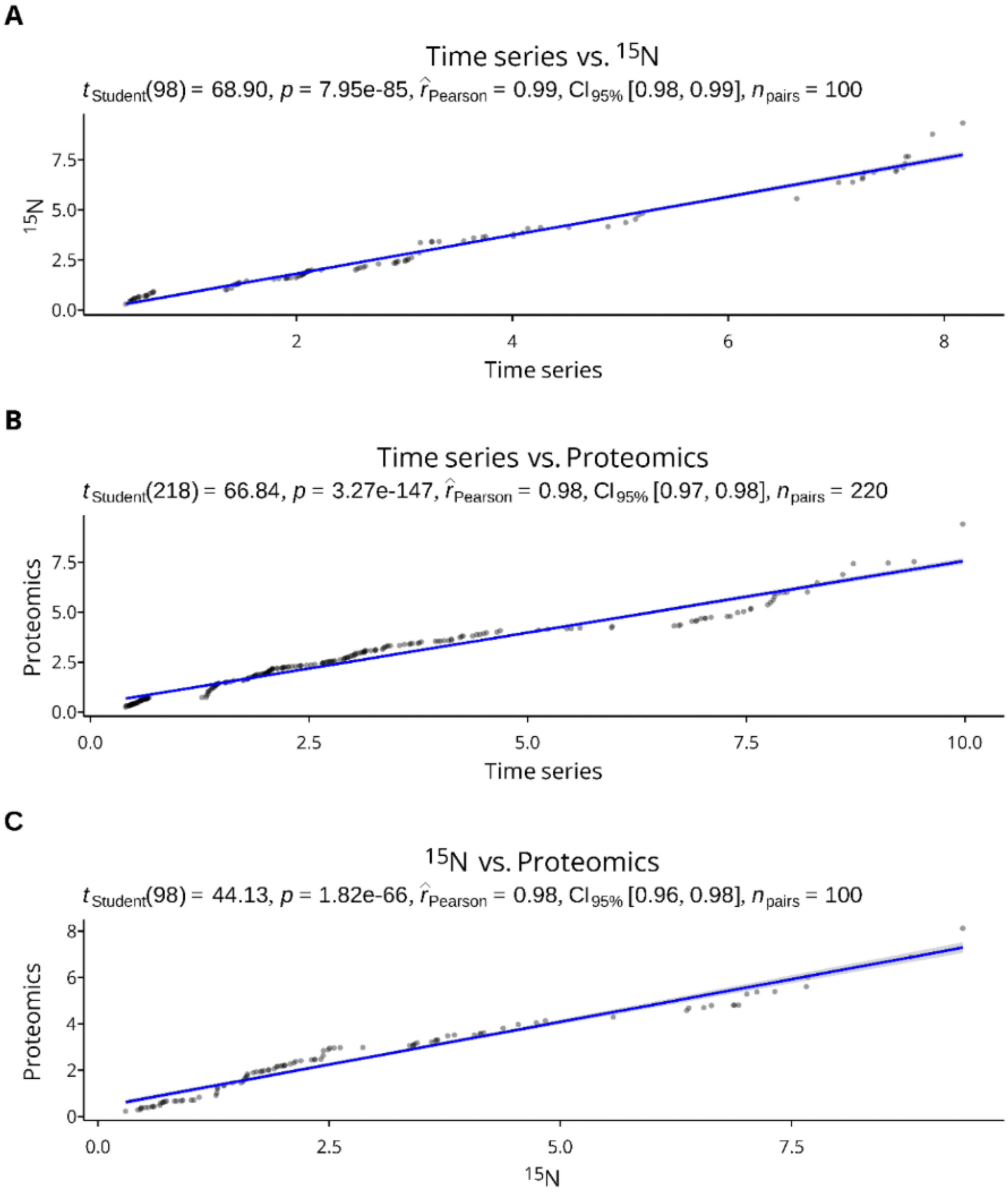


**Supplemental Figure S5**

Correlation matrices of projected leaf area between all 3 performed experiments. A) Correlation matrix between time series and $^{15}N$ (r = 0.99); B) Correlation matrix between time series and Proteomics (r = 0.98); C) Correlation matrix between $^{15}N$ and proteomics (r = 0.98). Correlations have been calculated using the Pearson correlation coefficient r.

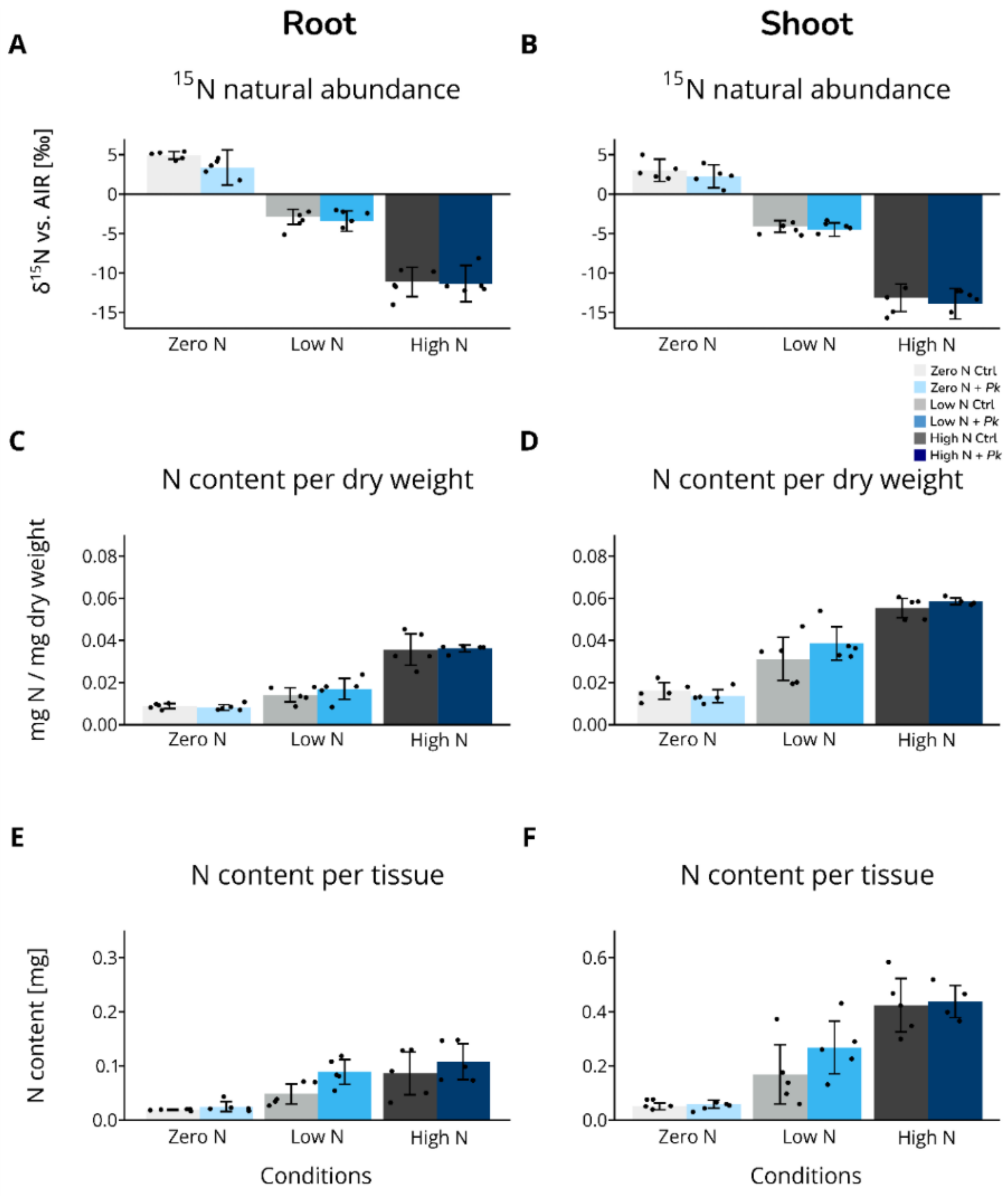


**Supplemental Figure S6**

Measurement of $^{15}N$ natural abundance in plant tissue at various N availabilities. Data presented are the mean $\delta^{15}N$ signatures vs. AIR (n = 5) given as deviation from the medium ($\delta^{15}N$ vs. AIR of medium is subtracted) ± standard deviation. Individual replicates are shown as dots on top of their respective bars. Means were compared using ANOVA followed by Tukey honestly significant difference (HSD) test.

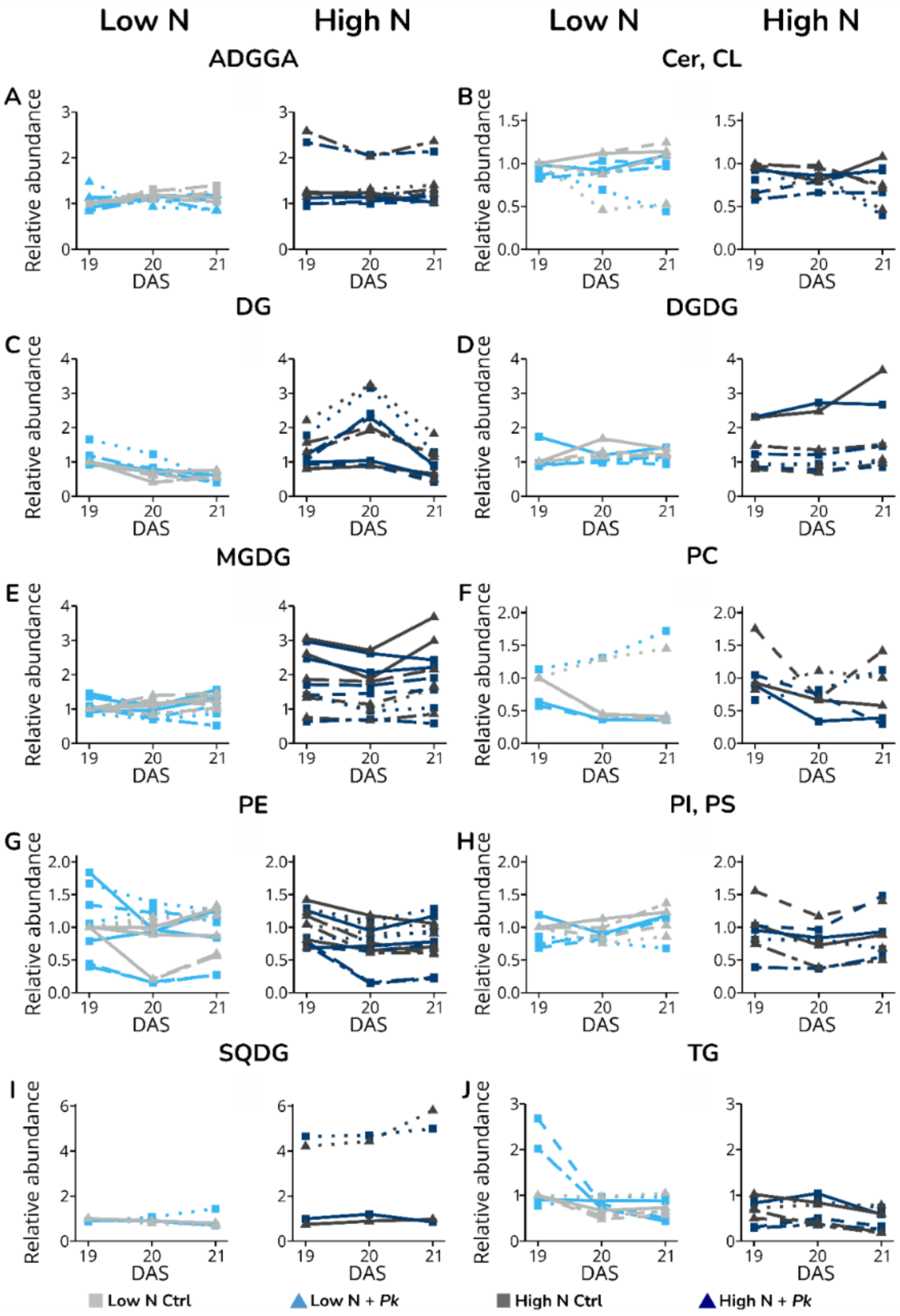


**Supplemental Figure S7**

Relative abundance of lipids with significant change in abundance in at least one time point in the comparison: Low-N + *Pk* vs. Low-N Control in roots. Lipids were split according to their classifications (ADGGA; Cer; DG, DGDG; LPC; MGDG; PC; PE; PI; SQDG; TG). Unique features are represented with different line types, with a maximum of 4 different features per plot. Relative abundances were calculated based on the abundances of features in low-N controls at 19 DAS (set to 1). Statistics can be found in the Supplemental table 2. Abbreviations: ADGGA, acyl diacylglycerol glucuronide; Cer, ceramides; DG, diacylglycerol; DGDG, digalactosyldiacylglycerol; LPC, lysophosphatidylcholine; MGDG, monogalactosyldiacylglycerol; PC, phosphatidylcholine; PE, phosphatidylethanolamine; PI, phosphatidylinositol; SQDG, sulfoquinovosyl diacylglycerol; TG, triacylglycerol.

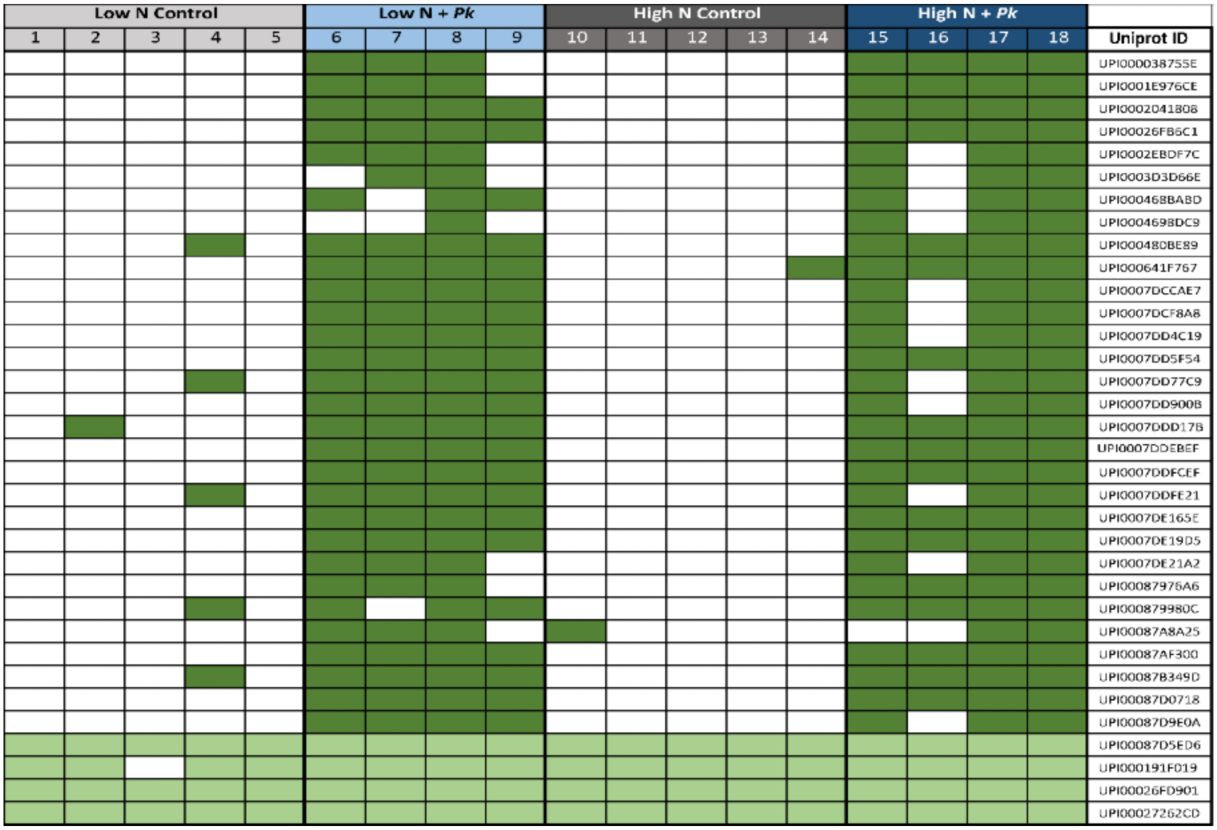


### Supplemental Figure S8

Presence or absence of *Pk*-derived proteins across individual samples. Dark green indicates detection of *Pk*-specific proteins; light green indicates conserved bacterial proteins present in all samples.

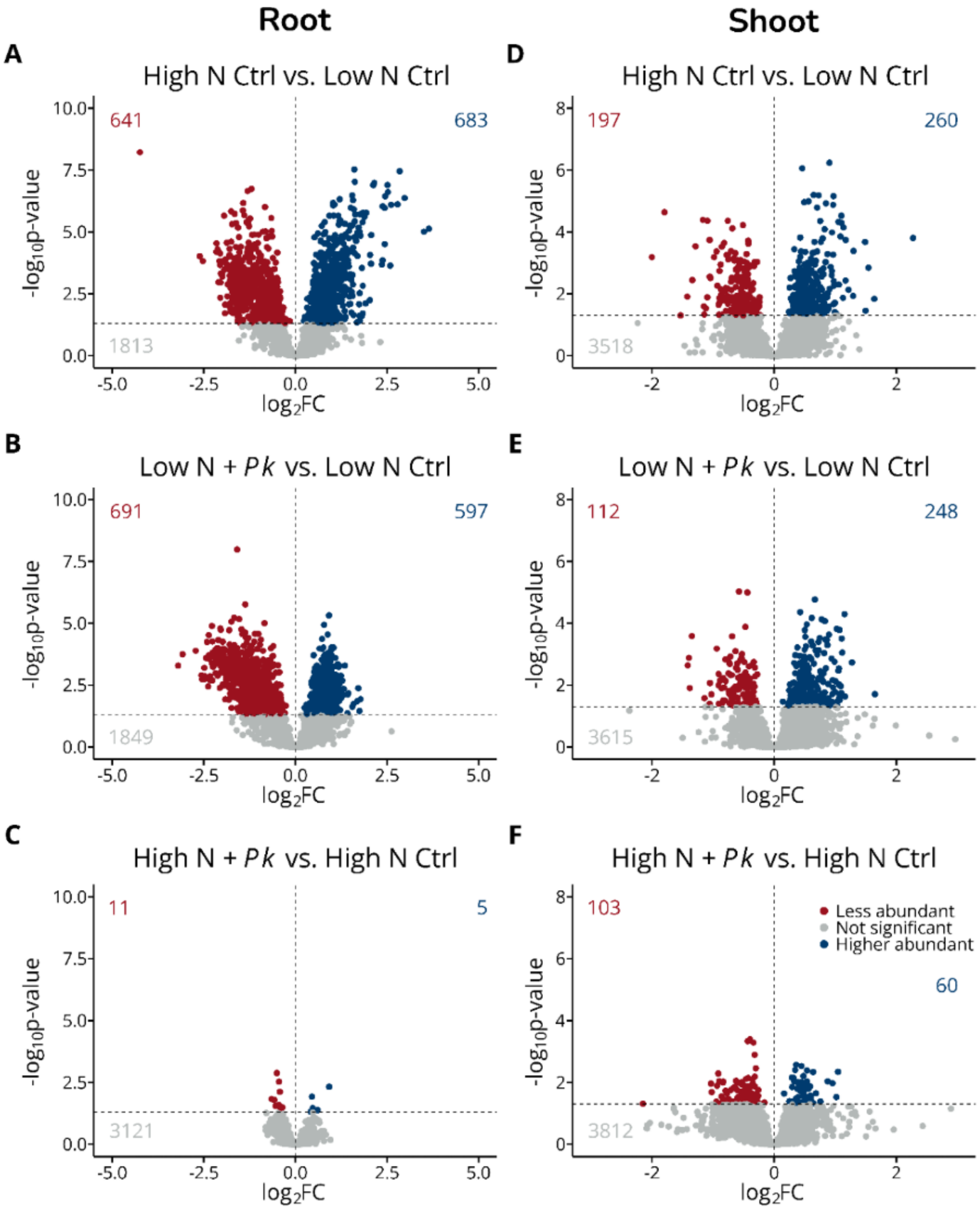


**Supplemental Figure S9**

A) Root volcano plot of the comparison: High-N Ctrl vs. Low-N Ctrl; B) Root volcano plot of the comparison: Low-N + *Pk* vs. Low-N Ctrl; C) Root volcano plot of the comparison: High-N + *Pk* vs. High-N Ctrl; D) Shoot volcano plot of the comparison: High-N Ctrl vs. Low-N Ctrl; E) Shoot volcano plot of the comparison: Low-N + *Pk* vs. Low-N Ctrl; F) Shoot volcano plot of the comparison:

High-N + *Pk* vs. High-N Ctrl; Data shown are 100% valid values of the respective data set. Volcano plots represent the -log10 transformed adjusted p values from Tukey's HSD with a p-value threshold set for α <0.05 and no threshold set for log2FC (= 0). Values in the plot represent the number of proteins in each group (red = less abundant, grey = not significant, blue = higher abundant; matching the colours of data points).

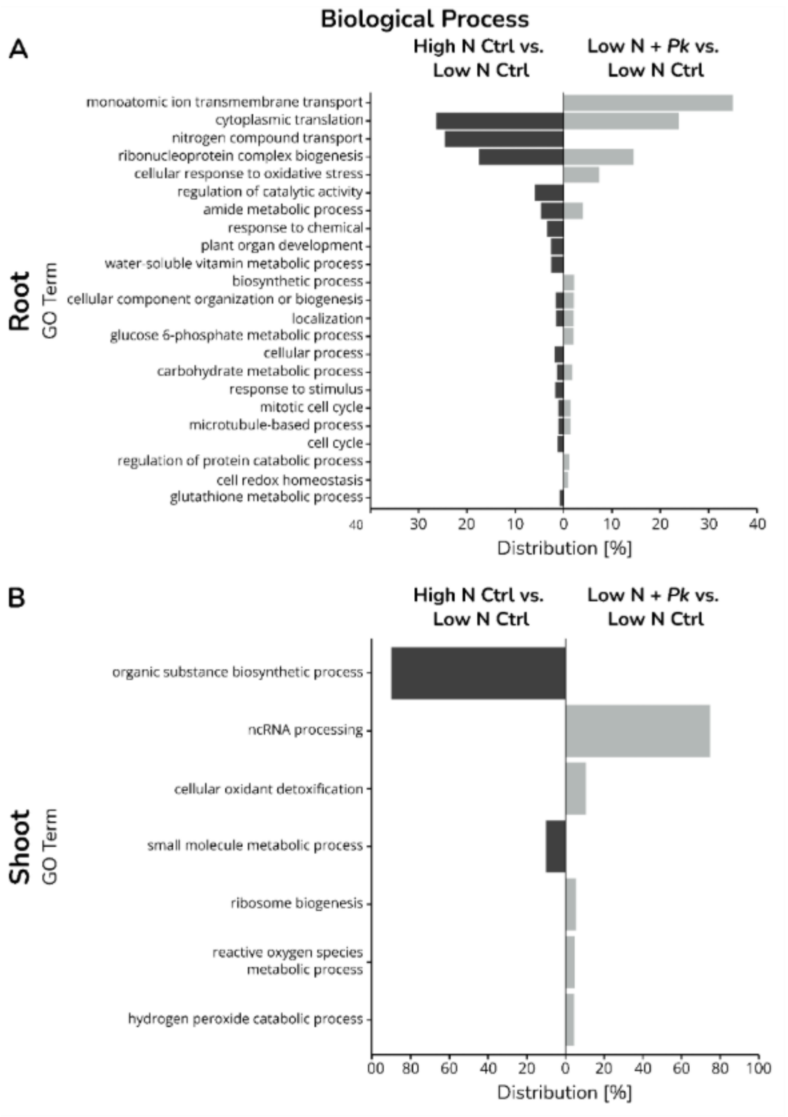


### Supplemental Figure S10

GO term analyses of Biological Processes. Only statistical significant proteins were imported into PantherDB statistical enrichment test (Mi et al., 2019), followed by REVIGO (Supek et al., 2011). Log10 (p-values) were used to calculate the distribution in percent (x-axis) of GO terms (y-axis). A) Root biological process GO-Terms; B) Shoot biological GO-Terms.

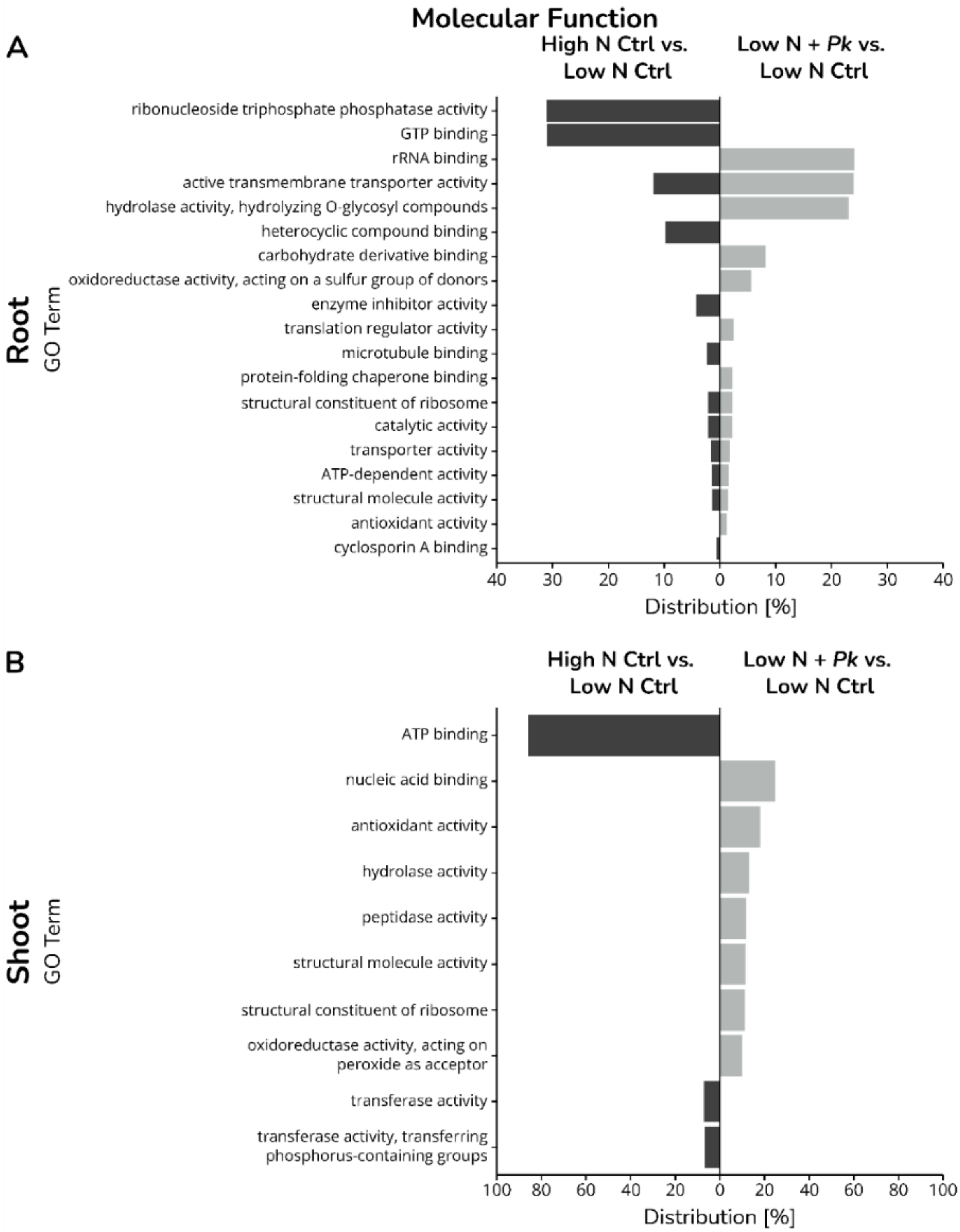


**Supplemental Figure S11**

GO term analyses of Molecular Functions. Only statistical significant proteins were imported into PantherDB statistical enrichment test (Mi et al., 2019), followed by REVIGO (Supek et al., 2011). Log10 (p-values) were used to calculate the distribution in percent (x-axis) of GO terms (y-axis). A) Root molecular function GO-Terms; B) Shoot molecular function GO-Terms.

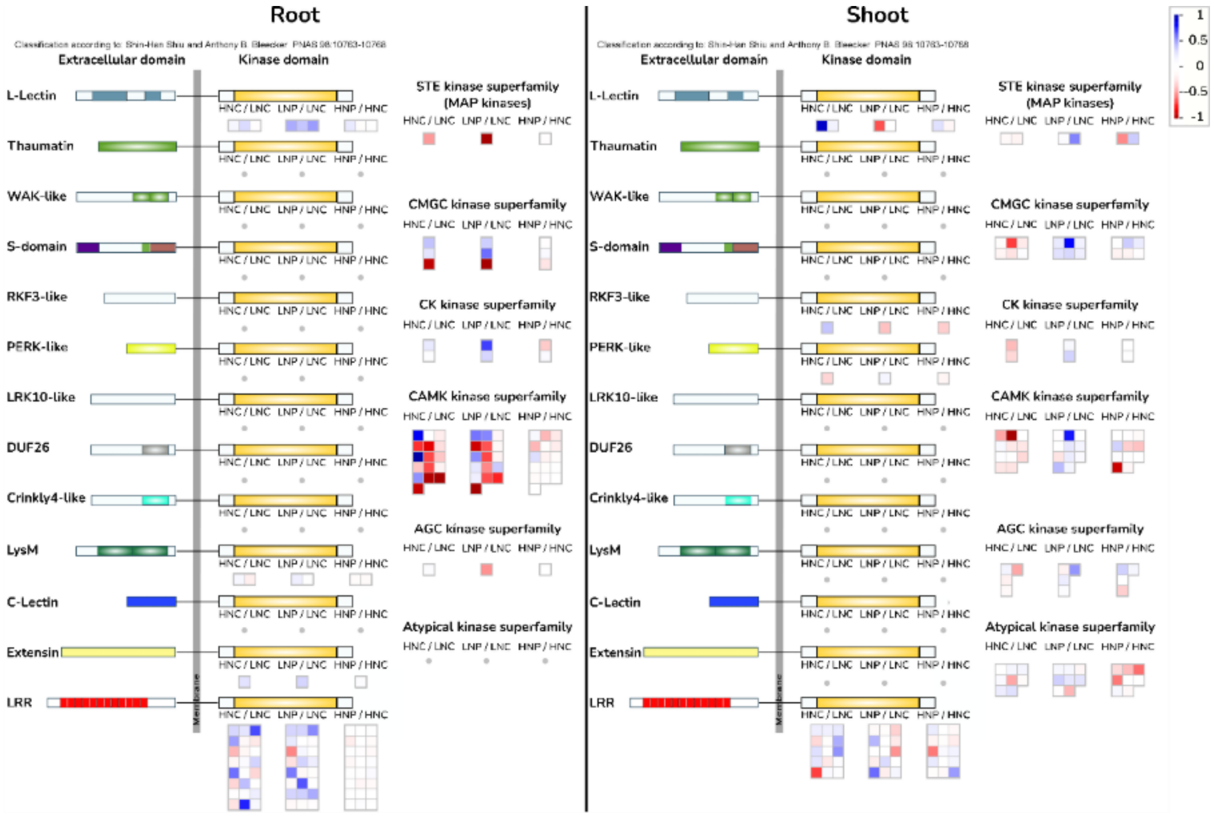


**Supplemental Figure S12**

Ratio of protein kinase abundance in roots and shoots, depicted as heatmap with blue indicating higher abundance and red indicating lower abundance.. Data represented are 100% vv. Mapped protein IDs, their log2FC and significances (indicated with bold value in log2FC column) are shown in supplemental table 2. MapMan version 3.5.1R2 (Thimm et al., 2004) was used. The following pathways were used and modified from Shiu and Bleecker, 2001: X4.1_Kinase_Families_R1.0. Mapping file was created using Mercator4 V2.0 (Schwacke et al., 2019). Comparisons are mentioned on top of each box: HNC / LNC = High-N Control vs. Low-N Control; LNP / LNC = Low-N + *Pk* vs. Low-N Control; HNP / HNC = High-N + *Pk* vs. High-N Control. The numerical values for this figure can be found in supplemental table 1.

### Supplemental Tables

Supplemental Table S1

Mapped proteins of the N-metabolism (Fig. 7), kinase families (Supplemental Fig. S12) and root lipid metabolism (Fig. 6). The column 'Mapped under' refers to the caption used in each pathway figure. Log2FC values in **bold** are significant different in their respective comparison. In case the proteins were not found in a tissue, log2FC were replaced with ‘x’, and ‘o’ if not mapped (shoot lipid metabolism). Statistics were performed using ANOVA with a confidence interval of 95% followed by post-hoc Tukey HSD.

| | | | **Root log2FC** | | | **Shoot log2FC** | | |
|---|---|---|---|---|---|---|---|---|
| | **Mapped under** | **Uniprot ID** | **HNC / LNC** | **LNP / LNC** | **HNP / HNC** | **HNC / LNC** | **LNP / LNC** | **HNP / HNC** |
| **N-metabolism (Fig. 7)** | AA | I1I0M5_BRADI | **-1.69** | **-1.41** | -0.19 | x | x | x |
| | | I1IDC0_BRADI | -0.70 | -0.70 | 0.48 | x | x | x |
| | AMT | I1IBN3_BRADI | **-1.56** | **-1.64** | -0.29 | x | x | x |
| | | I1IZJ3_BRADI | x | x | x | -0.13 | 0.10 | **-0.84** |
| | AS | A0A2K2D9P1_BRADI | x | x | x | **0.87** | **0.83** | -0.01 |
| | | I1H6K4_BRADI | -0.88 | **-1.78** | -0.22 | x | x | x |
| | GDH | I1GM99_BRADI | 0.26 | **0.89** | -0.09 | **-0.41** | -0.07 | 0.00 |
| | | I1J084_BRADI | **0.49** | -0.05 | 0.03 | 0.00 | -0.22 | 0.16 |
| | GOGAT | I1GRK9_BRADI | 0.29 | **0.78** | 0.13 | 0.25 | 0.28 | -0.23 |
| | | I1HQF1_BRADI | -0.41 | -0.23 | 0.07 | **0.27** | **0.43** | -0.10 |
| | GS1 | I1H7X9_BRADI | 0.11 | **-1.00** | -0.21 | **0.40** | **0.30** | 0.14 |
| | | I1IFI7_BRADI | **0.83** | -0.07 | 0.05 | -0.30 | -0.04 | 0.22 |
| | GS2 | I1J2T4_BRADI | 0.34 | -0.37 | 0.16 | 0.13 | 0.19 | -0.15 |
| | NiR | I1IEV4_BRADI | 0.40 | **1.04** | 0.33 | **1.08** | **1.07** | -0.15 |

| | | | | | | | | |
|---|---|---|---|---|---|---|---|---|
| | Nitrogen regulation | A0A0Q3MUB2_BRADI | **0.67** | **0.71** | 0.13 | -0.06 | 0.12 | 0.00 |
| | NR | A0A0Q3FHC7_BRADI | **-1.07** | -0.49 | 0.10 | **0.38** | 0.15 | 0.07 |
| | NRT | I1GPH0_BRADI | x | x | x | 0.01 | 0.08 | -0.02 |
| | | I1H1D1_BRADI | 0.70 | 0.83 | -0.14 | -0.27 | -0.15 | 0.36 |
| | | I1HBQ0_BRADI | x | x | x | 0.20 | 0.31 | -0.04 |
| | | I1I5R4_BRADI | **-1.34** | **-1.14** | -0.27 | -0.04 | -0.02 | -0.16 |
| | | I1I6K3_BRADI | **-1.24** | **-1.47** | -0.19 | x | x | x |
| | | I1IAY0_BRADI | x | x | x | 0.11 | 0.38 | -0.10 |
| | | I1IB70_BRADI | **-4.24** | 0.13 | -0.15 | x | x | x |
| **Kinase families (Supplemental Figure S12)** | AGC kinase superfamily | I1I858_BRADI | 0.08 | **-0.53** | -0.03 | x | x | x |
| | | a0a0q3hgt3_BRADI | x | x | x | 0.47 | 0.52 | 0.15 |
| | | I1GP76_BRADI | x | x | x | -0.16 | -0.18 | 0.34 |
| | | I1HHS0_BRADI | x | x | x | 0.19 | 0.22 | -0.03 |
| | | I1IWY7_BRADI | x | x | x | 0.13 | 0.09 | -0.35 |
| | Atypical kinase families | a0a0q3im64_bradi | x | x | x | -0.12 | 0.35 | -0.59 |
| | | i1gl81_bradi | x | x | x | -0.28 | **-0.43** | -0.13 |
| | | i1hf02_bradi | x | x | x | -0.21 | -0.08 | 0.14 |
| | | i1hkj6_bradi | x | x | x | -0.15 | 0.14 | -0.41 |

| | | | | | | | | |
|---|---|---|---|---|---|---|---|---|
| | | i1hm71_bradi | x | x | x | -0.16 | -0.14 | -0.61 |
| | | i1iav0_bradi | x | x | x | 0.47 | 0.25 | -0.03 |
| | | i1imj5_bradi | x | x | x | 0.02 | 0.15 | -0.31 |
| | | i1j2e3_bradi | x | x | x | 0.10 | 0.26 | -0.02 |
| | CAMK kinase superfamily | i1gkm6_bradi | **-0.73** | **-0.91** | -0.43 | 0.24 | 0.25 | -0.35 |
| | | i1glt7_bradi | -0.38 | -0.17 | -0.10 | x | x | x |
| | | i1gmg8_bradi | -0.39 | 0.25 | -0.09 | x | x | x |
| | | i1gmz0_bradi | **-0.86** | -0.66 | -0.08 | x | x | x |
| | | i1gu02_bradi | **-0.68** | **-0.55** | -0.09 | 0.32 | 0.12 | 0.00 |
| | | i1gx02_bradi | **-0.93** | **-1.11** | -0.03 | x | x | x |
| | | i1h269_bradi | -0.18 | 0.37 | -0.06 | x | x | x |
| | | i1hi64_bradi | **0.83** | **0.59** | 0.13 | x | x | x |
| | | i1hny1_bradi | 0.02 | 0.56 | -0.43 | 0.02 | 0.04 | -0.11 |
| | | i1hpi9_bradi | **0.53** | 0.20 | -0.11 | 0.25 | 0.42 | -0.92 |
| | | i1i692_bradi | -0.29 | -0.13 | -0.22 | -0.08 | -0.26 | 0.08 |
| | | i1iae4_bradi | **1.05** | 0.54 | 0.29 | 0.05 | 0.22 | -0.05 |
| | | i1in49_bradi | **-0.97** | **-0.67** | -0.07 | x | x | x |
| | | i1ix12_bradi | -0.22 | -0.15 | -0.24 | -0.13 | 0.03 | 0.34 |
| | | i1ixt7_bradi | **-0.68** | **-0.69** | -0.18 | x | x | x |
| | | i1j0z2_bradi | **-1.16** | -0.75 | -0.03 | x | x | x |
| | | I1GQB4_BRADI | x | x | x | 0.24 | 0.16 | -0.40 |
| | | I1H626_BRADI | x | x | x | **1.64** | 0.77 | -0.07 |
| | | I1H6L6_BRADI | x | x | x | **0.47** | -0.07 | -0.02 |
| | | I1ITM2_BRADI | x | x | x | 0.26 | 0.16 | -0.14 |
| | CK kinase superfamily | I1HFR4_BRADI | 0.23 | 0.70 | -0.37 | 0.42 | 0.13 | 0.02 |

| | | | | | | | | |
|---|---|---|---|---|---|---|---|---|
| | | I1HQU4_BRADI | 0.08 | 0.32 | 0.11 | 0.32 | 0.34 | 0.06 |
| | CMGC kinase superfamily | I1GNX9_BRADI | 0.39 | 0.35 | -0.07 | 0.23 | 0.19 | 0.22 |
| | | I1GUL3_BRADI | 0.26 | 0.61 | 0.12 | x | x | x |
| | | I1H4A1_BRADI | **-0.95** | **-1.05** | -0.25 | -0.08 | 0.21 | -0.07 |
| | | I1HAG7_BRADI | x | x | x | 0.09 | 0.23 | 0.04 |
| | | I1HGU1_BRADI | x | x | x | 0.16 | 0.27 | -0.17 |
| | | I1HIE2_BRADI | x | x | x | 0.27 | **0.32** | -0.17 |
| | | I1I8I8_BRADI | x | x | x | **0.70** | **0.81** | 0.36 |
| | Extensin | A0A0Q3IUM4_BRADI | 0.27 | 0.33 | 0.09 | x | x | x |
| | L-Lectin | a0a0q3kpd2_bradi | 0.15 | 0.46 | 0.24 | **-0.93** | **-0.67** | 0.28 |
| | | a0a2k2chp2_bradi | 0.31 | 0.38 | -0.09 | -0.18 | -0.03 | -0.15 |
| | | i1i8j5_bradi | -0.06 | 0.50 | -0.05 | x | x | x |
| | LRR | a0a0q3eyz3_bradi | **0.74** | 0.35 | -0.32 | **0.31** | **0.43** | -0.17 |
| | | a0a0q3gl83_bradi | **0.50** | **0.32** | 0.10 | -0.53 | -0.53 | 0.18 |
| | | a0a0q3gu79_bradi | -0.05 | 0.01 | -0.19 | -0.08 | 0.02 | 0.15 |
| | | a0a0q3kbu7_bradi | **0.91** | 0.88 | -0.05 | -0.26 | -0.21 | 0.23 |
| | | a0a0q3kl20_bradi | **0.48** | **0.49** | -0.09 | x | x | x |
| | | a0a0q3kre6_bradi | -0.20 | -0.05 | -0.03 | 0.09 | -0.01 | -0.08 |

| | | | | | | | | |
|---|---|---|---|---|---|---|---|---|
| | | a0a2k2cik2_bradi | -0.12 | 0.50 | -0.01 | x | x | x |
| | | a0a2k2ctf7_bradi | -0.06 | **-0.19** | -0.08 | x | x | x |
| | | a0a2k2d9s6_bradi | -0.48 | -0.14 | 0.17 | 0.24 | 0.31 | 0.00 |
| | | i1gp23_bradi | **1.02** | **0.82** | -0.06 | -0.20 | -0.09 | -0.02 |
| | | i1gwg0_bradi | 1.13 | -0.14 | 0.11 | x | x | x |
| | | i1h403_bradi | -0.36 | -0.10 | -0.14 | x | x | x |
| | | i1h6f1_bradi | **-0.65** | **-0.84** | 0.06 | x | x | x |
| | | i1ha83_bradi | 0.16 | 0.50 | 0.03 | x | x | x |
| | | i1hbq9_bradi | -0.23 | 0.22 | -0.10 | x | x | x |
| | | i1hde7_bradi | -0.01 | 0.04 | -0.19 | x | x | x |
| | | i1hib3_bradi | **-0.49** | -0.18 | 0.05 | x | x | x |
| | | i1hp99_bradi | -0.23 | -0.42 | -0.20 | -0.29 | 0.01 | -0.59 |
| | | i1hsu8_bradi | 0.61 | **0.99** | -0.22 | x | x | x |
| | | i1id77_bradi | 0.14 | **0.52** | 0.03 | **0.70** | 0.61 | -0.08 |
| | | i1ilv2_bradi | 0.31 | **0.70** | 0.01 | x | x | x |
| | | i1ip60_bradi | -0.41 | -0.18 | 0.02 | x | x | x |
| | | i1iva7_bradi | 0.39 | 0.49 | -0.04 | -0.21 | -0.12 | 0.22 |
| | | i1iz22_bradi | 0.20 | -0.10 | -0.05 | -0.12 | -0.21 | -0.12 |
| | | I1GMT9_BRADI | x | x | x | -0.07 | 0.15 | **0.51** |
| | | I1GR39_BRADI | x | x | x | -0.46 | -0.23 | -0.22 |
| | | I1GZ29_BRADI | x | x | x | -0.24 | -0.34 | -0.13 |
| | | I1IW92_BRADI | x | x | x | 0.06 | -0.01 | 0.13 |
| | LysM | A0A0Q3FRN3_BRADI | -0.20 | 0.05 | -0.12 | x | x | x |
| | | A0A0Q3I004_BRADI | 0.14 | 0.20 | -0.10 | x | x | x |

| | | | | | | | | |
|---|---|---|---|---|---|---|---|---|
| | MAP kinases | I1I5N8_BRADI | -0.51 | **-1.33** | -0.07 | x | x | x |
| | | I1H0Y7_BRADI | x | x | x | 0.17 | **0.56** | **0.36** |
| | PERK-like | A0A0Q3MRU1_BRADI | x | x | x | **0.33** | 0.16 | **-0.16** |
| | RKF3-like | A0A0Q3G6V2_BRADI | x | x | x | -0.38 | -0.40 | -0.36 |
| **Lipid metabolism (Fig. 6)** | acetyl-COA | A0A0Q3K0H7_BRADI | 0.26 | 0.16 | 0.16 | o | o | o |
| | | A0A0Q3M1M0_BRADI | 0.40 | 0.41 | 0.04 | o | o | o |
| | | I1GQ11_BRADI | 0.22 | -0.16 | 0.33 | o | o | o |
| | | I1H338_BRADI | **-0.92** | -0.55 | -0.28 | o | o | o |
| | | I1HLK9_BRADI | 0.29 | 0.17 | 0.10 | o | o | o |
| | | I1HQ21_BRADI | **-1.75** | **-1.13** | -0.09 | o | o | o |
| | | I1I6C4_BRADI | **-1.14** | -0.65 | -0.31 | o | o | o |
| | | I1IA43_BRADI | **-0.93** | **-1.20** | -0.15 | o | o | o |
| | | I1IQ05_BRADI | 0.39 | -0.03 | -0.22 | o | o | o |
| | | I1ITV5_BRADI | **-0.63** | **-0.68** | -0.13 | o | o | o |
| | | I1IVQ0_BRADI | 0.48 | -0.20 | 0.23 | o | o | o |
| | acetyl-CoA carboxylation | A0A0Q3JE04_BRADI | **-1.32** | **-1.01** | 0.02 | o | o | o |
| | | I1IWF2_BRADI | **-1.67** | **-1.99** | -0.22 | o | o | o |

| | | | | | | | | |
|---|---|---|---|---|---|---|---|---|
| | beta oxidation | A0A0Q3JRJ2_BRADI | **-0.94** | **-1.67** | -0.34 | o | o | o |
| | | I1HLG5_BRADI | **0.41** | 0.06 | -0.05 | o | o | o |
| | | I1HZH4_BRADI | -0.35 | -0.43 | -0.35 | o | o | o |
| | | I1I3F1_BRADI | -0.44 | **-0.87** | -0.23 | o | o | o |
| | | I1I4F7_BRADI | -0.10 | **-0.60** | -0.08 | o | o | o |
| | | I1IDX9_BRADI | **-0.54** | **-1.28** | -0.10 | o | o | o |
| | | I1IPG4_BRADI | -0.20 | **-0.73** | -0.27 | o | o | o |
| | desaturation | I1HN08_BRADI | -0.04 | 0.21 | -0.12 | o | o | o |
| | | I1HSR7_BRADI | **-1.77** | **-1.63** | 0.23 | o | o | o |
| | | I1HUM5_BRADI | 0.18 | 0.08 | -0.19 | o | o | o |
| | | I1HVL8_BRADI | 0.51 | 0.17 | -0.13 | o | o | o |
| | | I1I5F5_BRADI | 0.41 | 0.15 | -0.25 | o | o | o |
| | FAE | A0A2K2D3W9_BRADI | -1.10 | **-1.87** | -0.11 | o | o | o |
| | galactolipid sulfolipid | I1HIN3_BRADI | 0.18 | -0.04 | -0.02 | o | o | o |
| | | I1I6H2_BRADI | **1.93** | 0.28 | -0.45 | o | o | o |
| | glyoxylate cycle | A0A0Q3F6L1_BRADI | **0.54** | 0.24 | 0.04 | o | o | o |
| | | A0A0Q3GPP4_BRADI | -0.10 | **-0.76** | -0.37 | o | o | o |
| | | I1H6A1_BRADI | -0.48 | -0.41 | -0.09 | o | o | o |

| | | | | | | | | |
|---|---|---|---|---|---|---|---|---|
| | | I1HA00_BRADI | 0.12 | -0.35 | -0.07 | o | o | o |
| | | I1HZ21_BRADI | -0.54 | **-1.09** | -0.43 | o | o | o |
| | | I1IG62_BRADI | **-0.26** | **-0.47** | -0.09 | o | o | o |
| | | I1IX27_BRADI | **0.74** | **0.83** | 0.13 | o | o | o |
| | | I1IZE5_BRADI | -0.02 | -0.13 | 0.19 | o | o | o |
| | lipid bodies | A0A0Q3SCX9_BRADI | **-1.95** | **-1.64** | -0.02 | o | o | o |
| | | A0A2K2D5E4_BRADI | **-0.99** | **-0.79** | -0.53 | o | o | o |
| | | A0A2K2DPB0_BRADI | **-1.16** | **-0.99** | -0.40 | o | o | o |
| | | I1HLR0_BRADI | **-0.73** | **-0.61** | **-0.50** | o | o | o |
| | | I1IEG7_BRADI | 0.10 | 0.26 | 0.03 | o | o | o |
| | lipid trafficking | I1GL63_BRADI | -0.32 | **-0.68** | -0.10 | o | o | o |
| | | I1H1G1_BRADI | **0.65** | 0.57 | -0.26 | o | o | o |
| | | I1I1B4_BRADI | **1.06** | **0.93** | -0.10 | o | o | o |
| | lipid transport | A0A2K2D1A1_BRADI | **-1.12** | **-1.25** | -0.27 | o | o | o |
| | | A0A2K2DFT3_BRADI | 0.08 | -0.20 | 0.06 | o | o | o |
| | mtFAS | A0A0Q3R9W8_BRADI | 0.29 | 0.17 | -0.23 | o | o | o |
| | | I1H2L1_BRADI | **1.78** | 0.51 | -0.20 | o | o | o |

| | | | | | | | | |
|---|---|---|---|---|---|---|---|---|
| | | I1H5F3_BRADI | **1.34** | 0.83 | -0.18 | o | o | o |
| | | I1H6F4_BRADI | 0.41 | -0.09 | 0.19 | o | o | o |
| | phosphatidate | A0A0Q3FNL6_BRADI | **-1.18** | **-0.94** | -0.02 | o | o | o |
| | | I1I6G2_BRADI | 0.13 | 0.88 | -0.22 | o | o | o |
| | phosphatidylcholine | A0A0Q3GTN4_BRADI | **0.88** | 0.59 | -0.08 | o | o | o |
| | | A0A2K2D745_BRADI | -0.04 | **0.73** | -0.11 | o | o | o |
| | | I1GQE0_BRADI | **-0.97** | **-0.81** | 0.21 | o | o | o |
| | | I1HGU5_BRADI | 0.32 | 0.03 | -0.14 | o | o | o |
| | | I1HRW0_BRADI | **1.24** | 0.53 | 0.01 | o | o | o |
| | | I1HXR8_BRADI | **-0.89** | -0.61 | -0.04 | o | o | o |
| | phosphatidylethanolamine | A0A0Q3N4N1_BRADI | -0.22 | **0.90** | 0.07 | o | o | o |
| | | A0A2K2CRK8_BRADI | 0.36 | **0.53** | -0.18 | o | o | o |
| | | I1J2C7_BRADI | 0.29 | 0.13 | 0.08 | o | o | o |
| | phospholipase activities | A0A0Q3FHI6_BRADI | **-1.01** | -0.41 | -0.08 | o | o | o |
| | | A0A0Q3GLT1_BRADI | **1.16** | -0.58 | -0.33 | o | o | o |
| | | I1H4V2_BRADI | **-1.12** | **-0.68** | -0.21 | o | o | o |
| | | I1HCG5_BRADI | **0.58** | 0.18 | 0.22 | o | o | o |

| | | | | | | | | |
|---|---|---|---|---|---|---|---|---|
| | | I1HVB6_BRADI | **0.78** | 0.21 | -0.21 | o | o | o |
| | | I1I799_BRADI | 0.08 | 0.40 | -0.46 | o | o | o |
| | | I1IKD0_BRADI | **-1.26** | **-1.33** | -0.32 | o | o | o |
| | phytosterol | A0A0Q3NTK1_BRADI | **1.20** | 0.68 | 0.50 | o | o | o |
| | | I1GV37_BRADI | **-1.55** | **-2.07** | -0.26 | o | o | o |
| | | I1IVM8_BRADI | 0.04 | -0.18 | **-0.39** | o | o | o |
| | plasma membrane lipid transfer | I1H6Z5_BRADI | -0.18 | -0.19 | -0.11 | o | o | o |
| | plastid lipid transfer | I1IU18_BRADI | -0.38 | **-1.14** | -0.19 | o | o | o |
| | ptFAS | A0A0Q3IJU7_BRADI | 0.47 | **0.83** | -0.12 | o | o | o |
| | | A0A2K2CXZ2_BRADI | -0.39 | -0.71 | -0.06 | o | o | o |
| | | I1GKM7_BRADI | 0.06 | 0.55 | -0.18 | o | o | o |
| | | I1GXN7_BRADI | 0.71 | **0.98** | 0.19 | o | o | o |
| | | I1H029_BRADI | 0.71 | -0.48 | -0.03 | o | o | o |
| | | I1I9A1_BRADI | 0.28 | 0.64 | -0.04 | o | o | o |
| | | I1I9U8_BRADI | -0.18 | -0.04 | -0.32 | o | o | o |
| | | I1IRJ8_BRADI | **-0.76** | **-0.81** | 0.05 | o | o | o |
| | | I1IWW9_BRADI | 0.10 | 0.21 | -0.31 | o | o | o |

| | | | | | | | | |
|---|---|---|---|---|---|---|---|---|
| | sphingolipid | I1HPK0_BRADI | 0.00 | -0.04 | 0.12 | o | o | o |
| | | I1IBQ1_BRADI | 0.08 | -0.26 | -0.40 | o | o | o |
| | triacylglyerol lipase activities | I1HLP4_BRADI | 0.42 | -0.72 | -0.46 | o | o | o |

**Supplemental Table S2**

Number of identified proteins in each data set. The "100% valid values (vv)" column indicates proteins detected in 100% of the samples. The "ANOVA significant" column represents the number of significant proteins according to ANOVA (Perseus Multi-sample test, Permutation-based FDR = 0.05). The "Low-N + *Pk* vs. Low-N Ctrl (significant) column" shows the number of significant proteins in this specific comparison (statistics performed in R using two-way ANOVA followed by post-hoc Tukey's HSD).

| | **Identified proteins** | **100% vv each group** | **ANOVA significant (p-value <0.05)** | **High-N Ctrl vs. Low-N Ctrl (Tukey's HSD FDR <0.05)** | **Low-N + *Pk* vs. Low-N Ctrl (Tukey's HSD FDR <0.05)** | **High-N + *Pk* vs. High-N Ctrl (Tukey's HSD FDR <0.05)** |
|---|---|---|---|---|---|---|
| **Root** | 5660 | 3137 | 2291 | 1324 | 436 | 16 |
| **Shoot** | 6656 | 3780 | 1515 | 457 | 360 | 163 |

### Supplemental Methods

#### Change of 15N/$^{14}$N Isotopic distribution in plant N content

For $^{15}$N measurements (Supplemental Fig. S3, harvest III), sixty plants were grown as described, with the addition of a zero N condition, in which the medium did not contain $NH_4NO_3$, to account for N available in the seeds.

Plant samples were freeze-dried and 0.5 – 1.5 mg dry plant material was packed in tin capsules for stable nitrogen isotope ($\delta^{15}$N) analyses. Samples were combusted at 1060 °C with excess oxygen in an elemental analyser (Flash2000, Thermo Fisher, USA) and measured online with a coupled isotope ratio mass spectrometer (DeltaV plus, Thermo Fisher; USA).

International and laboratory standards (HZM (Holzmaar sediment), Acetanilide, Alanine, Histidine and Serin) were measured together with the samples. Calibration of laboratory standards and scale-normalisation of $\delta^{15}N$ raw values is based upon the international reference standards IAEA-N-2 ($\delta^{15}N$ = 20.3‰), IAEA-N-1 ($\delta^{15}N$ = 0.4‰) and USGS25 ($d^{15}N$ = -30.4‰).

Isotope results are reported as δ-values in ‰ according to the equation:

$\delta = R_S / R_{St} - 1$.

$R_S$ is the isotope ratio ($^{15}N/^{14}N$) of the sample and $R_{St}$ of the respective standard. δ-values for nitrogen are normalized to AIR (atmospheric nitrogen) scale (Coplen 2011). The average standard deviation of replicate measurements of standards was <0.20‰ for $\delta^{15}N$.
